# Natural variation in host defense strategies impacts both host and pathogen fitness

**DOI:** 10.1101/2022.09.22.509093

**Authors:** Alaina C. Pfenning-Butterworth, Rachel E. Vetter, Jessica L. Hite

## Abstract

1. Animals ranging from mosquitoes to humans often vary their feeding behavior when infected or merely exposed to pathogens. For example, some individuals drastically *reduce* their food intake (‘illness-mediated anorexia’) while others *increase* food intake (‘hyperphagia’). While these so-called ‘sickness behaviors’ are well documented, their functional consequences remain poorly resolved.
2. Here, we examine links between natural genetic variation in susceptibility to infection, feeding behaviors, multiple traits of the host, and within-host pathogen production. Using a zooplankton host (*Daphnia dentifera*) and a fungal pathogen (*Metschnikowia bicuspidata*) as a case study, we show that genotypic and dose-dependent variation in feeding behaviors are associated with both resistance and tolerance mechanisms.
3. In one genotype, immune-mediated anorexia was associated with increased tolerance to infection; unlike other genotypes, these individuals did not upregulate phenoloxidase activity, but lived longer, had the highest overall fecundity, and produced higher pathogen loads, despite their reduced growth rates and resultant smaller body sizes. In these hosts, peak parasite load remained unchanged, suggesting a tolerance mechanism that offset fecundity costs.
4. In other genotypes, feeding behaviors followed either a flat or hump-shaped pattern with pathogen dose, exhibiting hyperphagia at intermediate doses and anorexia at higher doses. In these cases, anorexia functioned primarily in resistance.
5. Our results suggest that infection-mediated changes in host feeding behavior — which are traditionally interpreted as immunopathology — may in fact serve as crucial components of host defense strategies. Moreover, these phenomena vary across host genotypes, and were associated with apparent trade-offs with another melanization component of immune defense. Together, these results underscore that while resistance and tolerance are typically viewed as alternative and fixed defense strategies, the immense genetic diversity for immune defense may result in more of a plastic spectrum spanning a gradient from resistance to tolerance.

## Introduction

Nutritional immunity is a powerful strategy at the core of the battle between host survival and pathogen proliferation. A host can prevent pathogen theft by sequestering essential nutrients, such as Mg, Fe, and Zn, or actively intoxicating pathogens with resource overload (Abu Kwaik and Bumann, 2013; Cunrath and Palmer, 2021), a phenomenon known as nutritional immunology or immunometabolism (Cotter et al., 2019; Pålsson-McDermott and O’Neill, 2020). Nutritional immunology frequently involves either an increase or decrease in food intake. These responses shape the within-host environment where pathogens proliferate and evolve and, in most cases, are considered a negative by-product of infection that directly contributes to host disease or immunopathology. However, mounting evidence reveals that reduced food intake or calorie restriction during infections can have numerous benefits ranging from improved immune function (Povey et al. 2014; Cotter et al. 2019) to reduced transmission (Rogers and Bates 2007; Rao et al. 2017). Beyond basic biology, resolving these mixed outcomes carries important practical implications for disease management in both natural and managed host populations.

A deeper understanding of the epidemiological and evolutionary implications of nutritional immunology (which here, includes associated changes in feeding behavior requires linking its impact on host and pathogen fitness, Graham et al. 2011; Hedrick 2017). Yet, the vast majority of empirical work on nutritional immunology and associated changes in feeding behavior has focused on the molecular and physiological drivers and little attention has been given to the effects of nutritional immunology on fitness costs beyond host survival (Ayres and Schneider 2009; Rao et al. 2017). In a recent review, only 19% of studies collected the data needed to quantify how nutritional immunology affects host or pathogen fitness (Hite et al. 2020a). Moreover, this work has generally taken a standard reductionist approach, focusing on immune stimulants (e.g., lipopolysaccharide) or single ‘LD-50’ doses. While, these studies have certainly advanced the field, results remain mixed and no broad-scale patterns have emerged: Inconsistent results across various host-pathogen systems, diverse pathogen genotypes, and differing infections routes challenge our understanding of the functional implications of this phenomenon (Hite et al. 2020a; Pike et al. 2019).

One reason for these variable responses involves the immense diversity of genetic-based traits that modulate immune responses in host populations. While the importance of host variation in immune defense is widely acknowledged, we know surprisingly little about how this variation affects functional outcomes, that is, host and pathogen traits. Moreover, this diversity is not well represented in the few inbred lab lines typical of model systems routinely used to study immune defense (Duffy et al., 2021; Graham, 2021). Changes in feeding behavior represent one axis in the defense arsenal a host enlists. The particular suite of defenses will vary, contingent on genetic variation in, for example, susceptibility to infection, current environmental conditions, or pleitotropic effects related to other life history traits (Moret and Schmid-Hempel, 2000; Schulenburg et al., 2009). Therefore, to gain a general understanding of the role that nutritional immunology plays in host-pathogen biology, we must account for the genetic diversity and environmental complexity present in the natural settings where hosts encounter pathogens.

As a first step in this endeavor, we use host genotypes sampled from nature to explicitly focus on the net effects of nutritional immunology and associated changes in feeding behaviors. Specifically, we examine links between changes in feeding behavior, host growth and reproduction, and parasite fitness (within-host spore load). Our work here was prompted by our identification that genotypes of the focal zooplankton host, *Daphnia dentifera*, vary in susceptibility to a virulent fungal pathogen, *Metschnikowia bicuspidata*, and illness-mediated feeding behaviors (Strauss et al., 2019). For instance, infection prevalence varied from 10-50% and some genotypes reduced food intake up to 78% (Strauss et al., 2019).

These findings and unique aspects of this host pathogen system (detailed below) allow us to address the following questions: Do more susceptible genotypes exhibit the strongest or weakest anorexia? Do these changes benefit the host, pathogen, both or neither? Of course, we recognize the causality dilemma inherent in these questions; hosts could have higher infection rates *because* they alter food intake. Addressing this dilemma, however, requires further investigation that is beyond the scope of our study. Here, we focus on understanding links between natural genotypic variation in these behaviors and traits. We hope that this first step will help uncover key patterns to guide future experiments in this and other host-pathogen systems with greater molecular resolution to identify more mechanistic and causal pathways.

The focal host-pathogen system is unique in that (1) transmission is relatively well understood (Ebert 2005; Stewart Merrill and Cáceres 2018; Hall et al. 2007) and can be readily re-created in the lab, (2) *Daphnia* are facultative parthenogens and typically reproduce asexually, and different isoclonal lines (hereafter: genotypes) often vary in key epidemiological traits like exposure (i.e. foraging rate) and per-spore susceptibility (Hall et al., 2010; Auld et al., 2013), (3) pathogen fitness can be easily approximated by quantifying spore yield, and (4) host fitness and other covarying life-history traits can be tracked over the entire course of infection.

Measuring these four intertwined parameters enables us to discriminate between resistance — the ability to control pathogen exposure, growth, or transmission — and tolerance — the reduction in infection-induced pathology or fitness costs (Schneider and Ayres, 2008; Read et al., 2008). Quantifying this collection of traits in the same study presents notable logistical challenges (Graham et al., 2011) and is understandably missing from most studies on nutritional immunology and associated changes in feeding behaviors.

Our results suggest that infection-mediated changes in host feeding behavior — which are traditionally interpreted as immunopathology — may in fact serve as crucial components of host defense strategies. Moreover, these phenomena vary across host genotypes, and were associated with apparent trade-offs with another melanization component of immune defense. These results also highlight that while resistance and tolerance are typically viewed as alternative and fixed defense strategies, there is enormous genetic diversity that results in variable defense strategies that likely result in more of a plastic spectrum spanning a gradient from resistance to tolerance.

### Study System

The *Daphnia - M. bicuspidata* system presents a rare opportunity to quantify links between two immune responses (phenoloxidase and anorexia/hyerphagia), how they jointly influence multiple traits of the host as well as pathogen fitness across pathogen exposure levels, and how these patterns vary across host genetic backgrounds. The focal hosts, *D. dentifera*, are key consumers in aquatic food webs throughout northern temperate lakes where they are hosts to numerous pathogens, including the virulent pathogen studied here, *M. bicuspidata*. Hosts become infected while filter feeding on phytoplankton. Upon successful infection, the fungal spores pierce through the host’s gut wall into the body cavity and avoid degradation by host haemocytes (Metschnikoff, 1884; Stewart Merrill and Cáceres, 2018). This ‘obligate-killer’ pathogen exhibits a parasitoid life history strategy that requires the death of its host for onward transmission. However, before the pathogen can kill the host and release spores into the water column where they infect new zooplankton hosts, it must escape several innate immune responses.

*Daphnia* have a relatively simple, albeit powerful, innate immune response. These responses include phagocytosis by haemocytes (Metschnikoff, 1884; Stewart Merrill and Cáceres, 2018), phenoloxidase (Mucklow and Ebert, 2003), and changes in immune-mediated feeding behavior (Hite et al., 2017; Strauss et al., 2019). In this study, we focus on the latter two immune responses. Dietary changes, including total calorie intake, can affect the abundance of phenoloxidase (Pauwels et al., 2010; Povey et al., 2014; González-Santoyo and Córdoba-Aguilar, 2012). Phenoloxidase is a key component in the molecular pathway that produces melanin, which attaches to pathogens to hinder their growth and reproduction (Cerenius and Söderhäll, 2004). *Daphnia* (and other invertebrates) increase active phenoloxidase (PO) abundance upon exposure to pathogens (Labbe and Little, 2009). Moreover, *Daphnia* genotypes vary in their phenoloxidase activity and this variation is negatively correlated with infection success (Mucklow and Ebert, 2003; Mucklow et al., 2004).

## Methods

### Combined Foraging and Infection Assay

We measured immune-mediated feeding behavior in adult females (6-day old) from four unique host genotypes across a gradient of three pathogen exposure levels (Hite et al., 2017; Strauss et al., 2019). All genotypes were chosen from existing cultures isolated from lakes in Michigan or Indiana (USA). Cultures were maintained in high nitrogen COMBO, artificial lake water media (Kilham and Herrera, 1998) and fed lab-cultured *Ankistrodesmus falcatus* (1mgC/L). This algae resembles the shape (needle-like) and size (40–50 µm long; 3–5 µm wide) of *M. bicuspidata* spores. Hosts were maintained at 22°C for at least three generations to standardize any maternal affects.

We cultured *M. bicuspidata in vivo* in a standard genotype of *D. dentifera*. We standardized all conditions known to affect the infection success of the pathogen, including temperature (Shocket, 2018), host age (Hite et al., 2017), algal quality and quantity (Hall et al., 2009), and spore age (Duffy and Hunsberger, 2019). Current evidence indicates that genetic differences among pathogen strains do not affect transmission success (Searle et al., 2015; Duffy and Sivars-Becker, 2007; Shaw and Duffy, 2021). These results suggest that hosts traits predominately shape the outcomes of this host-pathogen interaction. All spores used in the infection assay were collected 21 days prior to the assay and counted with a hemocytometer.

We reared cohorts of neonates from each host genotype for five days (under a 16:8 light:dark photoperiod at 22*°*C). Then, individuals were isolated in vials (volume V = 10 ml) containing 1mgC/L of *A. falcatus*. We inoculated thirty replicates of each genotype at each density of fungal spores: 0, 150, or 300 spores/ml. We treated control vials identically to spore treatments, except that we did not add hosts. That is, controls contained algae and pathogen spores but no hosts to consume the algae or spores. To ensure that algae and spores remained suspended throughout the assay, we gently inverted all vials every 30 minutes. We conducted the entire assay (set-up to take-down) in the dark to prevent algal growth or any spurious spikes in algal fluorescence. Host were exposed to spores for 24 hours, but we measured feeding rates after seven hours (based on Hite et. al., unpublished data).

At the end of the seven-hour feeding rate assay, we collected subsamples from each vial to measure *in vivo* fluorescence following Sarnelle and Wilson 2008; Hite et al. 2020b. In brief, this method compares the fluorescence of algae in vials with hosts vs. the fluorescence of algae in the host-free (consumer-free) controls. We measured algal fluorescence using narrow-band fluorometry (Tecan, Maennedorf, Switzlerand) with 200 µL in each well (each with two technical replicates) of a black 96-well plate (14-245-197A Thermo Fisher Scientific No. 7605). For extended technical details see Hite et al. 2020b.

We calculated the feeding rate of individual hosts, *f* following Sarnelle and Wilson (2008), by solving for the change in fluorescence, *F*:

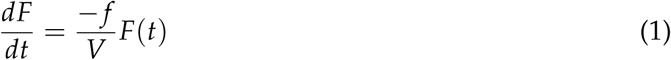

Where *F*_*t*_ is the food remaining (the mean algal fluorescence of the sample at time, *t*), *F*_0_ is the initial amount of food (the mean algal fluorescence of the corresponding plate-specific animal-free controls at time *t* = 0), *V* is the volume of media (10 mL), and *t* is the length of the assay. Solving for *f*, then:

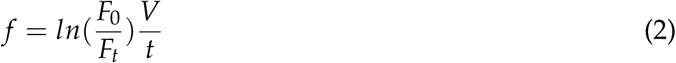

We omitted any negative feeding rates (since these represent technical errors and provide a more conservative estimate of immune-mediated anorexia since strong negative feeding rates could bias the results), individuals that died during the assay, and animals identified as male (male and female *Daphnia* have different feeding rates, Hite et al. 2017).

### Life History Table

To examine the relationship between immune-mediated feeding behaviors, phenoloxidase, and host susceptibility, we measured the proportion of hosts that became infected and the survivorship of infected hosts relative to uninfected hosts. Then, to determine how immune-mediated feeding behaviors and phenoloxidase affect the realized fitness of both hosts and pathogens, we quantified host growth, fecundity, and total pathogen load. We maintained individuals at 22*°*C in vials 15mL of COMBO for fourteen days post-exposure (or death, whichever came first). Individuals were moved to fresh media, fed (1mgC/L for the first seven days and 2mgC/L for the rest), and checked for offspring every other day.

We visually diagnosed terminal infections with a dissecting microscope, measured final body size (at 5x magnification), and collected individuals to quantify spore loads using flow cytometry. We used a DxP10 flow cytometer (Cytek) equipped with a BD FACSort system (Becton Dickinson Biosciences, San Jose, CA, USA). To isolate mature transmission ready spores from algae, animal debris, or immature spores (Stewart Merrill and Cáceres, 2018), we used custom gates based on fluorescence forward scatter (FSC) and side scatter (SSC) with 488nm and 561nm lasers and fluorescent beads as standards (SPHERO, Accucount Fluorescent Particles, 7.0-8.0 um) at a ratio of 12:1 for each individual’s spore solution (1 animal in 300 µl of COMBO). We then verified flow cytometry spore counts by randomly selecting five individuals from each genotype and spore exposure level and manually counting spores using a hemocytometer (for Standard, R^2^ = 0.91; patterns for all other genotypes were comparable - see Appendix).

### Phenoloxidase Assay

We quantified phenoloxidase (PO) activity in the hemolymph (following Mucklow and Ebert 2003) from a subset of 239 (66%) animals. We collected hemolymph by pricking individual *D. dentifera* in the heart (26-gauge needle, Monject). The hemolymph of five–seven individuals was pooled to reach a final volume of 2µL and added to 150µL of PBS buffer on ice (0.15M NaCl, 10mM N*a*_2_HPO_4_·2H_2_0, pH 7.5). Next, we transferred 50µL of the hemolymph-PBS solution to 225µL of 20mM L-Dopa (with duplicate technical replicates). We measured the absorbance of each sample at 475nm immediately and at every hour interval for 4.5 hours (Tecan, Maennedorf, Switzerland). PO activity was calculated as the increase in absorbance after 4.5 hours (absorbance at 4.5 hours - absorbance at 0 hours) corrected by changes in the control (PBS and L-Dopa only). PO active units = (corrected change * 1000) / number of individuals in the sample (Mucklow and Ebert 2003).

### Statistical Analysis

All analyses were carried out in R statistical software. We used generalized linear models (GLMs) with host genotype (1, 2, 3, 4), initial host size, pathogen dose (0, 150, 300), infection status (control, exposed but uninfected, and exposed and infected), and their interaction as fixed effects. Frequency of infection was analyzed with binomial errors and log-link function. Because feeding rates are log transformed, they were always analyzed with Gaussian distributed errors and either identity or log link functions. For survival, we conducted both time-censored analyses and GLMs with longevity (time until death) as the response variable with Gaussian distributed errors and either identity or link functions. Both results were qualitatively similar (see Appendix). We present logevity in the main results because relative to the survival curves they are easier to decipher (in our opinion). We ran saturated and reduced models and used Akaike information criteria (AICc) for model selection with limited sample size (package: MuMIn). We then examined the selected models using the residual diagnostics, summary, and accessed significance using Wald chi-square statistics (package: car) following Fox 2011. For *post hoc* analyses, we used least square means with Tukey’s correction for multiple comparisons. The full code and statistical analyses are available together with the data on GitHub.

## Results

### Susceptibility

As expected, the focal genotypes varied in susceptibility to infection (Fig. 2). The ‘Standard’ and ‘Midland 281’ genotypes were the most and least susceptible genotypes, respectively. Relative to ‘Standard’ and ‘Midland 281’, the other two genotypes exhibited intermediate levels of susceptibility. The dose-dependent effects on infection frequency were variable across genotypes and was not a statistically significant predictor of infection frequency, even after accounting for differences in host body size (Table 1). Though not statistically significant, for two gentoypes (‘Standard’ and ‘Midland 252’), infection frequency tended to decrease with higher pathogen exposure levels; whereas, with ‘Midland 281’ and ‘A45’ infection frequency tended to increase at higher doses (Fig. 2).

**Figure 1.**
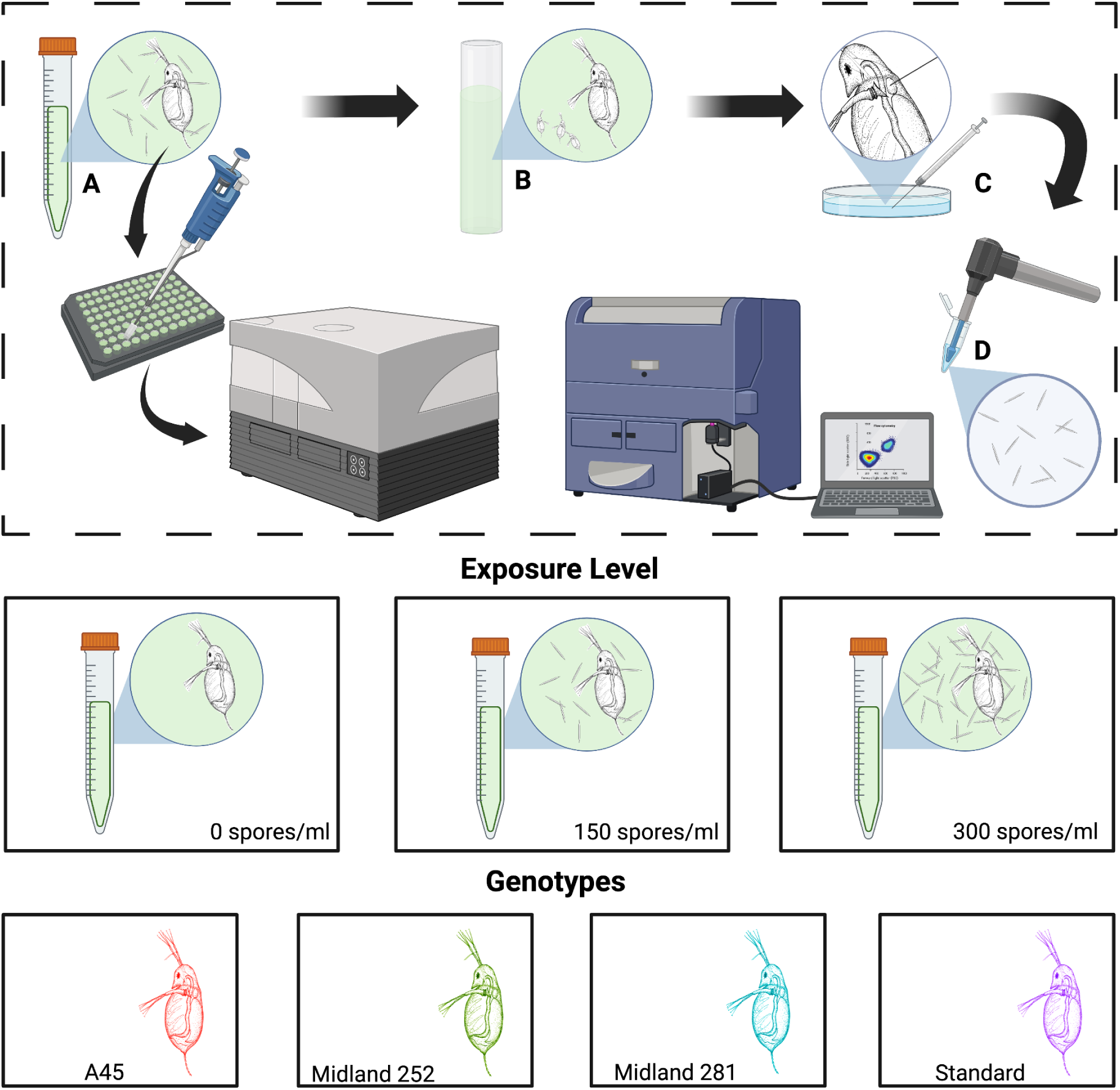
Experimental design to study links between susceptibility to infection and the net effects of nutritional immunology and associated changes in feeding behavior. We used the *Daphnia dentifera*-*Metschnikowia bicuspidata* host-pathogen system as a case study. These facultative parthenogenetic hosts produce asexual broods of females throughout most of the year, which enables a unique opportunity to examine both genetic and environmental variation, namely pathogen exposure levels (indicated in the figure by the density of small needle-like spores). **(A)** In a joint feeding-rate infection assay, we tracked changed in immune-mediated feeding behaviors and susceptibility to infection. **(B)** We then examined several life history traits of the hosts (day of first reproduction, number offspring produced each day) with a life table assay. **(C)** We then measured the final size of each individual, collected phenoloxidase (pooled 5 randomly-selected individuals within a genotype and spore exposure treatment), and **(D)** used flow cytometry to quantify spore yield from the remaining infected individuals. To standardize the collection of spore yield, the experiment was terminated on 10 days post exposure (when the first individual died). This limits our ability to quantify effects of infection on host longevity. However, a central goal of this study was to examine effects on pathogen load at a biologically-reasonable time point during infection.

**Figure 2.**
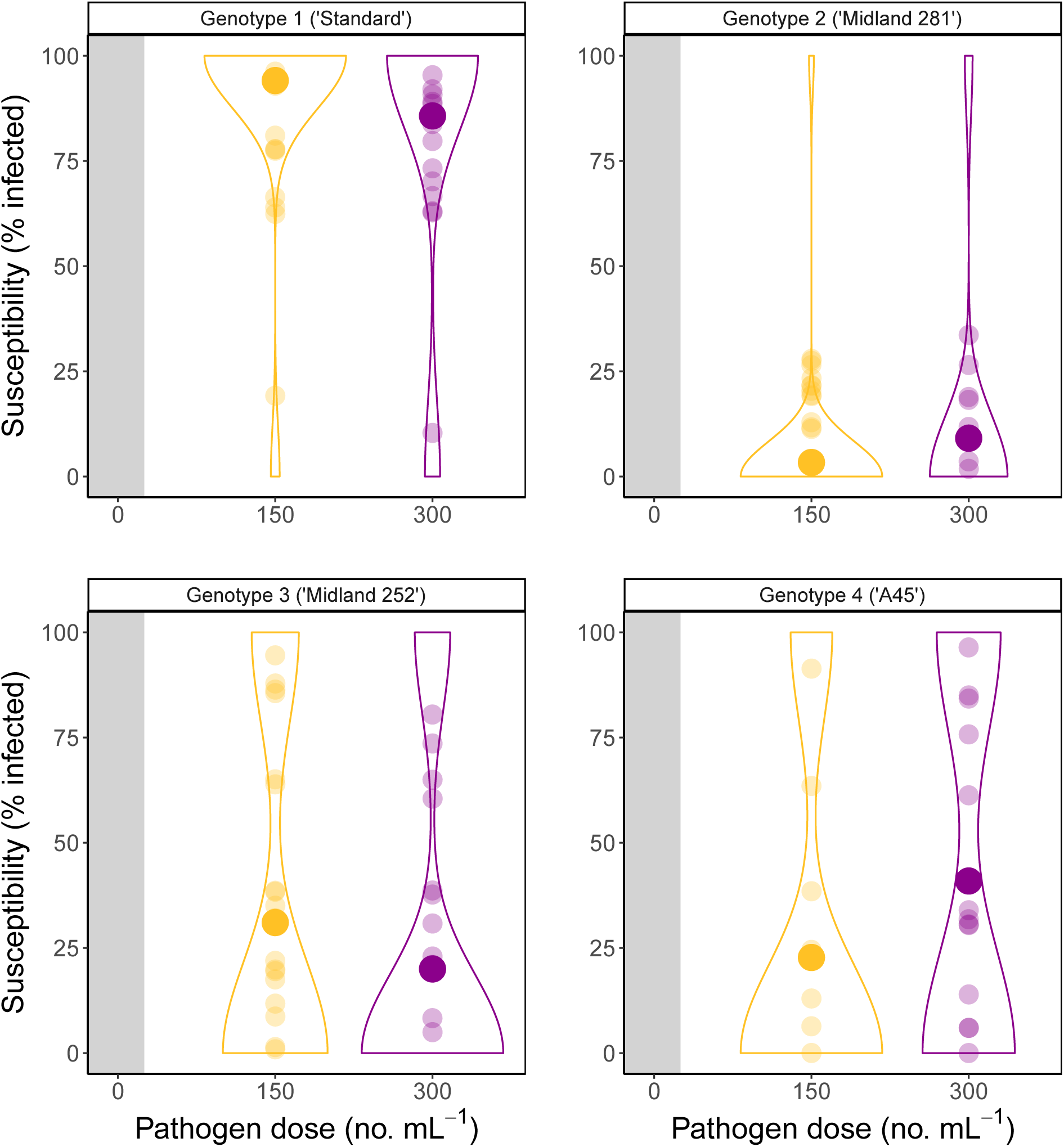
Natural variation in host susceptibility to infection as a function of pathogen exposure levels. Small symbols indicate replicates and large symbols indicate the mean.

**TABLE 1.** Factors influencing host susceptibility to infection (% infected). See Methods for further details on infection assays and statistical analysis.

| **Term** | **Wald $\chi^2$** | **df** | **Pr( > $\chi^2$)** |
| --- | --- | --- | --- |
| Genotype | 32.15 | 3 | **4.8x10<sup>-7</sup>** |
| Pathogen exposure level | 0.023 | 1 | 0.880 |

Bold indicates significance values ( $p < 0.05$ ).

### Immune defenses

#### Feeding behaviors

We observed pronounced phenotypic variation in the defense mechanisms of focal host genotypes. For example, analysis of total food consumption, a metric commonly used to quantify illness-mediated anorexia and hyperphagia, showed that two genotypes altered their feeding behavior in a dose-dependent manner and two genotypes did not (Fig. 3). For the ‘Standard’ genotype, infected (and exposed) individuals decreased food intake by up to 42%, depending on spore dose (Fig. 3). On the other hand, infected (or exposed) individuals from the ‘Midland 281’ genotype, either *increased* food intake (hyperphagia), by approximately 90% at intermediate pathogen doses or decreased food intake by 32% at higher doses (Fig. 3). Notice that ‘Midland 281’ (which had the lowest susceptibility, Fig. 2) consumed much higher food (and thus pathogen spores) relative to the other three genotypes (compare axis scaling in Fig. 3). For the other two genotypes (with intermediate levels of susceptibility), infection had minor effects on feeding rates (Fig. 3). Within each genotype, infected individuals consumed less food relative to their uninfected counterparts (all p-values *<* 0.001), but there was no difference in food intake between infected and exposed individuals (all p-values *>* 0.05), even after Tukey correction and accounting for size-dependent variation (see Appendix for full results). Together, these results indicate that for *Daphnia*, illness-mediated anorexia and hyperphagia are both genotype- and dose-dependent (Table 2).

**Figure 3.**
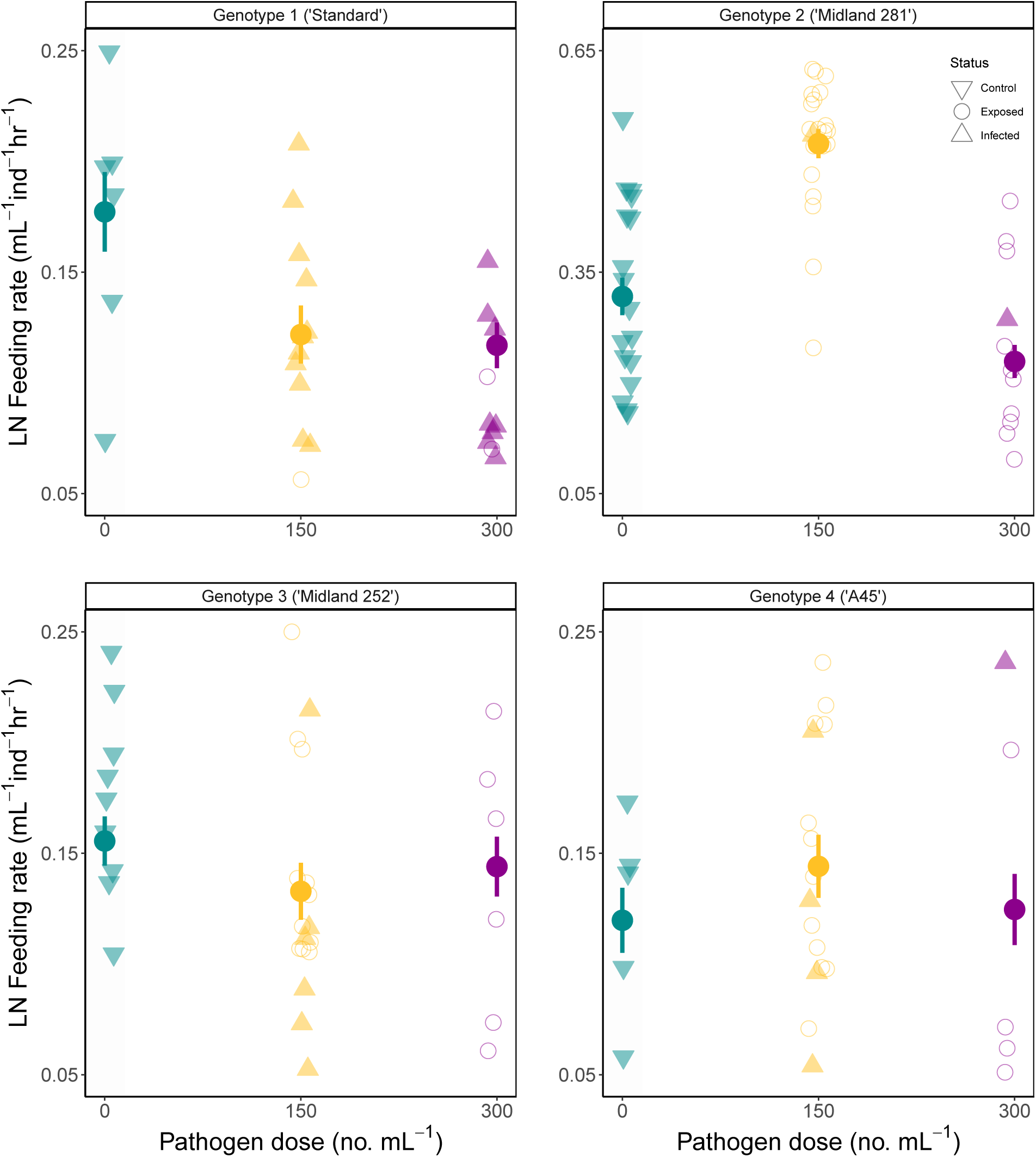
Genotypic and spore-dependent changes in feeding behavior as a function of pathogen infection or exposure. Filled circles are means *±* standard errors with individual data points indicating hosts that were either never exposed to pathogens (downward triangles), exposed but remained uninfected (open circles), or exposed and infected (upward triangles).

**TABLE 2.** Factors influencing host feeding behavior. See Methods for further details on feeding rate assay and statistical analysis.

| **Term** | **Wald $\chi^2$** | **df** | **Pr( > $\chi^2$)** |
| --- | --- | --- | --- |
| Genotype | 219.45 | 3 | **2.6x10<sup>-47</sup>** |
| Initial body size (mm<sup>2</sup>) | 29.69 | 1 | **5.0x10<sup>-08</sup>** |
| Infection status | 90.69 | 2 | **2.0x10<sup>-20</sup>** |
| Pathogen exposure level | 78.06 | 1 | **9.9x10<sup>-19</sup>** |

Bold indicates significance values ( $p < 0.05$ ).

#### Phenoloxidase

Similar to feeding behaviors, we observed dose- and genotype-dependent differences in active phenoloxidase levels (Fig. 4, Table 3). Across all genotypes, phenoloxidase levels significantly increased at the highest exposure levels, but not at the intermediate dose. For ‘Standard’, changes in phenoloxidase levels were slight, increasing a mere 4%, at the highest spore dose. However, for the other three genotypes, phenoloxidase increased by up to 41%. While the number of infected individuals included in these samples varied, this variation did not significantly affect these patterns (Table 3). Together, these results indicate that similar to changes in feeding behavior, phenoloxidase is both genotype- and dose-dependent. Moreover, for at least two genotypes (‘Standard’ and ‘Midland 281’), defense appears to rely on *either* changes in feeding behavior *or* phenoloxidase, which suggests that these two immune mechanisms do not work in tandem. Whether these differences are a cause or consequence of differences in immune defense is, however, difficult to tease apart.

**Figure 4.**
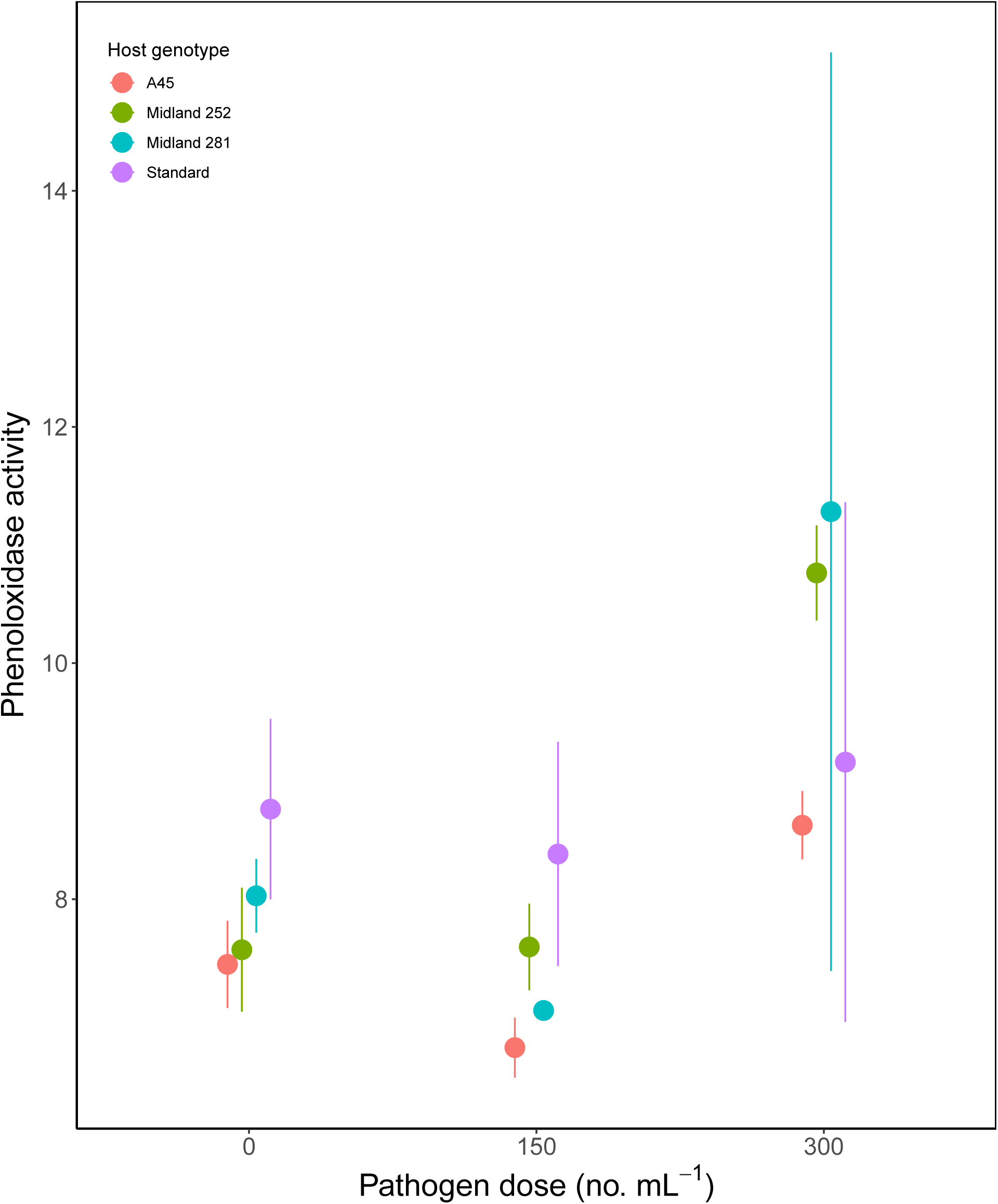
Genotypic and spore-dependent variation in phenoloxidase activity (PO). We collected hemolymph from a subset of 239 (66%) of *D. dentifera* and pooled the hemolymph of five-seven individuals to reach sufficient concentrations. Data are means *±* standard error.

**TABLE 3.** Factors influencing active phenoloxidase (PO) in hosts. See Methods for further details on infection assays and PO collection.

| **Term** | **Wald $\chi^2$** | **df** | **Pr( > $\chi^2$)** |
| --- | --- | --- | --- |
| Genotype | 15.42 | 3 | 0.001 |
| Pathogen exposure level | 53.64 | 2 | **2.2x10<sup>-12</sup>** |

Bold indicates significance values ( $p < 0.05$ ).

### Net Outcomes

#### Linking host susceptibility to host and pathogen traits

Across all genotypes, susceptibility to infection and spore yield were positively correlated (r = 0.893, t = 22.39, *p <* 0.001, 95 %, CI = (0.852, 0.923), Fig. 5*A*). In other words, more susceptible genotypes tended to produce more spores and therefore, have higher spore yields (and transmission potential), regardless of pathogen exposure levels.

**Figure 5.**
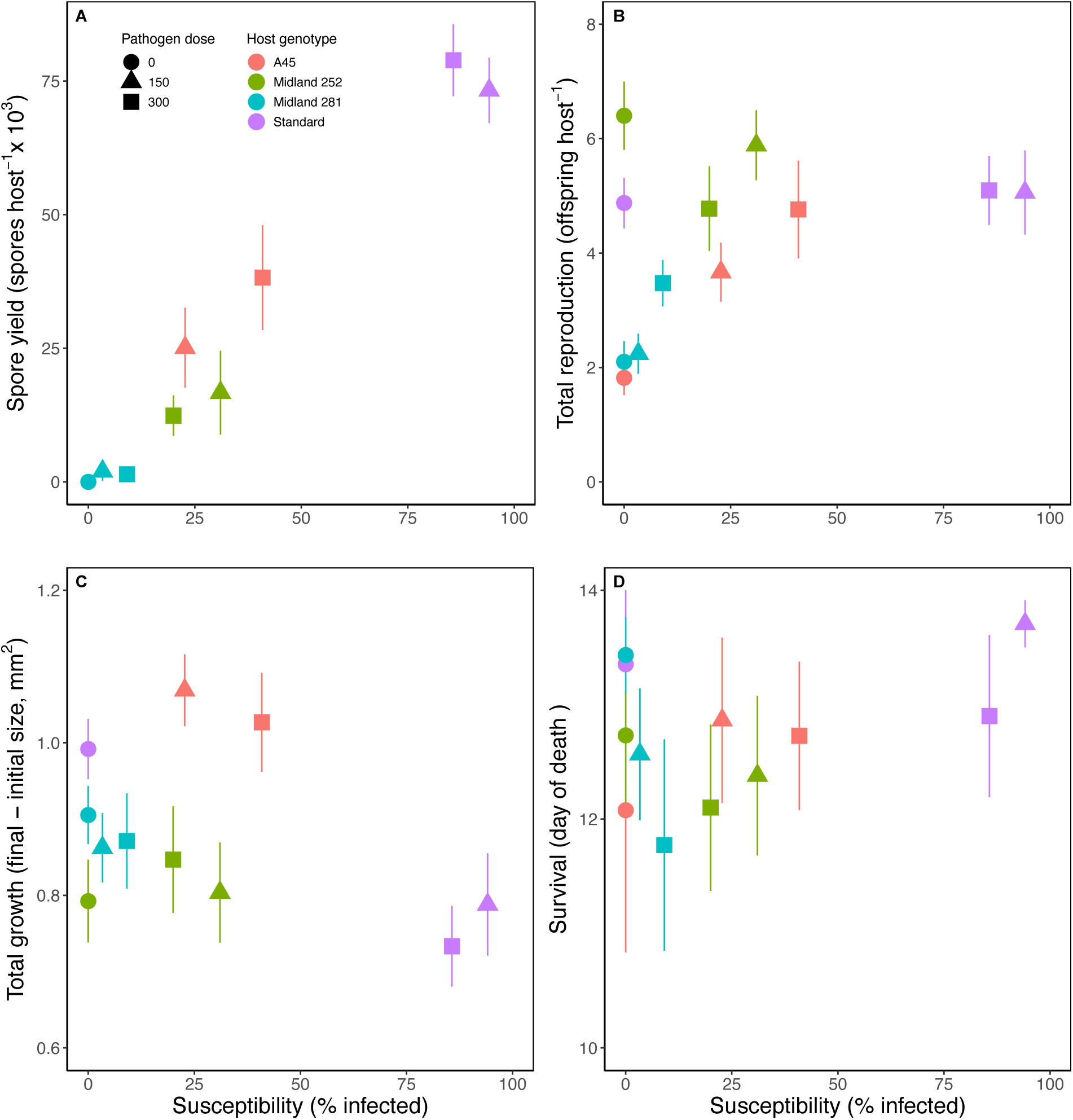
Connecting variation in susceptibility to infection to pathogen fitness and host life history traits. Focal host genotypes vary in **(A)** spore yield, a key metric of pathogen fitness, resulting in a positive relationship between susceptibility to infection and pathogen fitness. In other words, more susceptible genotypes also yielded higher spore yields. **(B)** Infection resulted in higher total reproduction over the course of the experiment, for all but one genotype (‘Midland 252’). **(C)** Infection had little net effect on host growth (except for the most susceptible genotype, ‘Standard’) or survival (at least over the course of the time-limited assay). Data are means *±* standard error.

How did this genotypic variation in susceptibility and spore yield, in turn, affect host fitness, growth, and survival? For the most susceptible genotype (‘Standard’), infection had no effect on host fitness (total reproduction; Fig. 5*B*). For the other genotypes, susceptibility to infection significantly influenced host fitness, even after accounting for differences in total growth, pathogen exposure level, and the interaction between genotype and exposure level (Table 4). For these genotypes, infection caused either a slight increase (‘A45’ and ‘Midland 281’) or a slight decrease in reproduction (‘Midland 252’). Infection also altered host growth (Table 4), especially for the most susceptible genotype, ‘Standard’ (Fig. 5*C*). Finally, susceptibility and pathogen exposure levels significantly reduced host survival (Fig. 5*D*), though there were no significant differences across host genotypes (Table 5). Patterns for these three host traits were qualitatively similar when we replaced ‘susceptibility’ with ‘infection status’ (see Appendix).

**TABLE 4.** Factors influencing host fitness (total offspring produced) and survival. See Methods for further details on the life history table and statistical analysis.

| **Term** | **Wald $\chi^2$** | **df** | **P value** |
| --- | --- | --- | --- |
| Susceptibility | 5.312 | 1 | **0.021** |
| Total Growth (mm<sup>2</sup>) | 21.31 | 1 | **3.9x10<sup>-6</sup>** |
| Genotype | 69.01 | 3 | **6.9x10<sup>-15</sup>** |
| Pathogen exposure level | 10.28 | 1 | **0.001** |
| Genotype · Path. exp. | 12.55 | 3 | **0.006** |

*Host growth (final – initial size)*
| **Term** | **Wald $\chi^2$** | **df** | **P value** |
| --- | --- | --- | --- |
| Susceptibility | 20.44 | 1 | **6.2x10<sup>-6</sup>** |
| Pathogen exposure level | 0.052 | 1 | 0.820 |
| Genotype | 35.57 | 3 | **9.2x10<sup>-8</sup>** |

*Host survival*
| **Term** | **Wald $\chi^2$** | **df** | **P value** |
| --- | --- | --- | --- |
| Susceptibility | 15.51 | 1 | **8.2x10<sup>-5</sup>** |
| Pathogen exposure level | 7.342 | 1 | **0.007** |
| Genotype | 1.108 | 3 | 0.775 |

Bold indicates significance values ( $p < 0.05$ ).

**TABLE 5a.** Factors influencing host susceptibility (% infected) and transmission potential (spore yield) **across all host genotypes. See Methods for further details on infection assays and statistical analysis.

| Term | Wald $\chi^2$ | df | Pr( > $\chi^2$) |
| --- | --- | --- | --- |
| Feeding rate | 12.27 | 1 | **4.6x10<sup>-4</sup>** |
| Initial body size (mm<sup>2</sup>) | 1.494 | 1 | 0.222 |

*Transmission potential (spore yield)*
| Term | Wald $\chi^2$ | df | Pr( > $\chi^2$) |
| --- | --- | --- | --- |
| Pathogen exposure level | 0.462 | 1 | 0.497 |
| Final body size (mm<sup>2</sup>) | 5.262 | 1 | **0.022** |

Bold indicates significance values (p < 0.05).

#### Linking feeding behaviors to host and pathogen traits

*Across all genotypes*, hosts that reduced food consumption (stronger anorexia) tended to have higher susceptibility (GLM: *χ*^2^ = 12.02, *p* < 0.001, Fig. 6*A*) and pathogen loads (spore yield) (GLM: *χ*^2^ = 18.98, *p <* 0.001, Fig. 6*B*). Notably, the most susceptible genotype (‘Standard’), which produced much higher spore yields, was also one of the smallest of the genotypes (Fig. 6*D*). This genotype, therefore, produced high spore yields and maintained reproductive levels similar to its uninfected counterparts (Fig. 5*B* and 6*C*), despite drastically reducing food intake (which should have also decreased ingestion of pathogen spores) and suffering reduced growth rates (Fig. 6*D*). Such pronounced differences in this genotype may help explain why, after accounting for genotypic differences, the effects of feeding rates and final host size (surface area, size^3^) on susceptibility were not statically significant (Table 5*b*).

**Figure 6.**
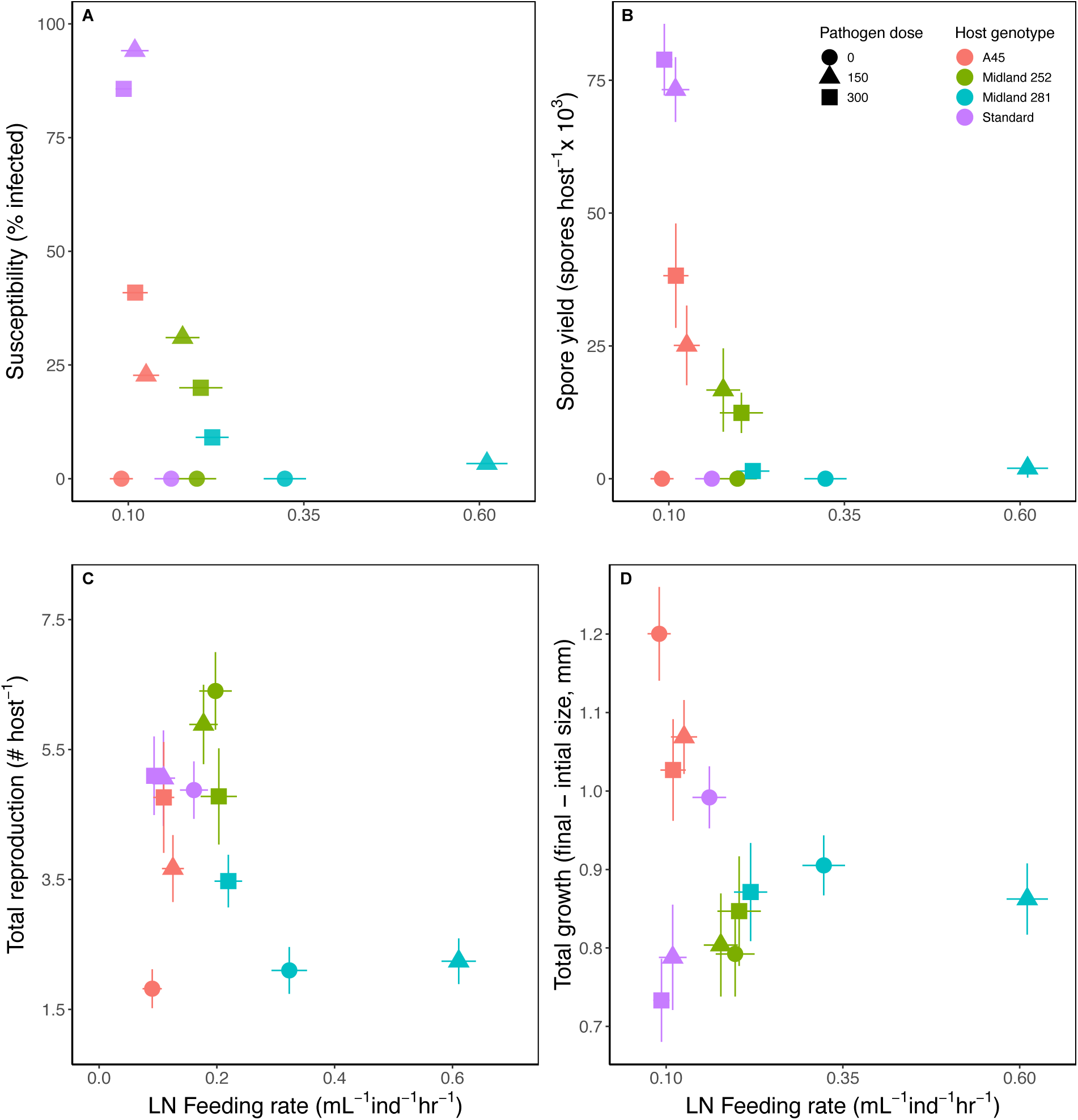
Connecting variation in feeding behavior to infection to key host and pathogen traits. **(A)** More susceptible genotypes tend to reduce food intake when infected and **(B)** produce more spores, a key metric of pathogen fitness. **(C-D)** Changes in feeding behavior attenuates the fecundity and growth costs of infection for the most susceptible and most tolerant genotypes, ‘Standard’ and ‘Midland 281’, respectively.

**TABLE 5b.** Factors influencing host susceptibility (% infected). and transmission potential (spore yield). See Methods for further details on infection assays and statistical analysis.

| Term | Wald $\chi^2$ | df | Pr( > $\chi^2$) |
| --- | --- | --- | --- |
| Feeding rate | 3.128 | 1 | 0.077 |
| Initial body size (mm<sup>2</sup>) | 3.622 | 1 | 0.057 |
| Pathogen exposure level | 0.103 | 1 | 0.748 |
| Genotype | 19.36 | 3 | **0.0002** |
| Feeding rate · Initial body size (mm<sup>2</sup>) | 3.581 | 1 | 0.058 |

*Transmission potential (spore yield)*
| Term | Wald $\chi^2$ | df | Pr( > $\chi^2$) |
| --- | --- | --- | --- |
| Genotype | 97.76 | 3 | **<2.2x10<sup>-16</sup>** |
| Pathogen exposure level | 51.57 | 2 | **6.34x10<sup>-12</sup>** |

Bold indicates significance values (p < 0.05).

#### The gestalt: Resistance, tolerance, and everything in between

Together, these results suggest that the defense response of ‘Standard’ (i.e., decreased food intake and no increase in phenoloxidase) helped offset the fecundity costs of infection, but at the expense of reduced growth. From the pathogen’s perspective, the tolerance of infection in ‘Standard’ led to high total pathogen loads, and therefore transmission potential, despite the smaller size of the host. However, for the least susceptible genotype, ‘Midland 281’, the host defenses (hyperphagia and low phenoloxidase at intermediate doses and anorexia and increased phenoloxidase at higher doses) functioned to: (a) suppress spore yield (anti-growth resistance) and (b) offset reproduction costs (at least at high doses, Fig. 5*B*; and growth Fig. 5*C*), while (c) tending to reduce host longevity (Fig. 5*D*). However, the qualitatively different results at the two pathogen doses hinders a clear mechanistic understanding of these phenomena. For the other two gentoypes with intermediate levels of susceptibility, defense strategies appear to be driven more by phenoloxidase than changes in feeding behaviors. This defense strategy resulted in intermediate levels of spore yield, which for ‘A45’ did not reduce fecundity, growth, or survival. For ‘Midland 252’ the higher production of phenoloxidase (relative to ‘A45’) may explain that genotype’s slight decrease in reproduction.

## Discussion

Here, we used host genotypes sampled from nature to explicitly examine links between changes in two focal immune responses, host fitness, and within-host parasite density. Tracking these intertwined parameters enabled us to formulate hypotheses regarding immune polymorphism and how natural selection shapes strategies for immune defense and pathogen transmission (Graham et al., 2011; Boots et al., 2009).

Our results suggest that shifts in feeding behaviors can function in both resistance and tolerance, depending on host genotype. For one genotype, illness-mediated anorexia was associated with increased host tolerance to infection, attenuating infection-associated declines in fitness; this came at the cost of reduced growth, but did not reduce the probability of infection or overall pathogen load (even after statistically accounting for smaller body size of hosts). The remaining three genotypes, however, did not exhibit immune-mediated anorexia. Instead, they upregulated phenoloxidase, resulting in lower fitness, but not reduced growth. For these genotypes, both pathogen load and the percentage of hosts that became infected were lower relative to ‘tolerant’ genotypes. Moreover, the magnitude of anorexia and hyperphagia were dose- and genotype-dependent, suggesting that such sickness behaviors are phenotypically plastic and possibly a consequence of genetic-based differences in defense strategies. Together, these results underscore that while resistance and tolerance are typically viewed as alternative and fixed defense strategies, the immense genetic diversity for immune defense may result in more of a plastic spectrum spanning a gradient from resistance to tolerance.

While patterns across all four genotypes were mixed, changes in food intake did help offset the fitness costs of infection and controlling pathogen populations within hosts (Fig. 6*B*). For instance, the ‘Standard’ genotype decreased food intake (Fig. 3), but did not increase in phenoloxidase (Fig. 4). This defense strategy offset the fecundity costs of infection (Fig. 5*B* and 6*C*), but at the expense of reduced growth (Fig. 5*C* and 6*D*). Through an evolutionary epidemiology lens (Graham et al., 2011), the ‘Standard’ genotype tolerates infection, leading to higher total pathogen loads, and therefore transmission potential, despite the smaller size of the host. Together, these results join others in demonstrating that anorexia can function as a tolerance mechanism (Ayres and Schneider, 2009; Cumnock et al., 2018), at least for one of the focal genotypes.

On the other hand, ‘Midland 281’, enlisted a different defense strategy depending on the pathogen dose in the environment. At intermediate doses, individuals consumed more food (hyperphagia) and phenoloxidase levels remained low (Fig. 4). At higher doses, individuals consumed less food (anorexia) and increased phenoloxidase. Together, these dose-dependent strategies helped suppress pathogen populations (anti-growth resistance; Fig. 6*B*) and offset reproductive costs (at least at high doses, Fig. 6*C*-*D*) but with a reduction in host longevity (Fig. 5*D*), at least over the time frame captured in these assays. Changes in feeding behavior are known to be sensitive and finely-tuned first line of defense; the magnitude of anorexia (i.e., the decline in food intake) is typically dose-dependent, increasing with parasite exposure, infection intensity, parasitemia, or immune challenge. Our analysis extends previous studies by demonstrating that this dose-dependence may depend, in part, on the host’s genetically-determined susceptibility to pathogens (Fig. 5).

Together, our analysis suggests that for the genotypes studied here, the most susceptible genotype (‘Standard’) exhibits stronger anorexia, which appears to help both the host (by attenuating the reproductive costs of infection) and the pathogen (via higher pathogen loads). For the least susceptible genotype (‘Midland 281’), both hyperphagia and mild anorexia combined with phenoloxidase prevent infection and suppress pathogen populations in the few hosts that became infected. Interestingly, this genotype had relatively low fitness when uninfected, but higher fitness at the higher pathogen loads, suggesting a stress response (*sensu* life history theory (Stearns, 1992)) and little costs incurred for these immune responses. Further, these results underscore that efforts to evaluate when and why host responses cause more harm to the host (immunopathology) or the pathogen, must consider the effects on multiple host traits and pathogen fitness, especially in light of the immense genetic diversity for host immune defenses (Mucklow et al., 2004; Graham et al., 2011; Duffy et al., 2021).

For the other two genotypes with intermediate levels of susceptibility, defense strategies may be driven more by phenoloxidase than changes in feeding behavior. Moreover, phenoloxidase activity was finely tuned to the pathogen dose in the environment (Fig. 4). In arthropods, a cascade of serine proteases activates prophenoloxidase (PPO) and the enzyme phenoloxidase, which, in turn, catalyzes the synthesis of melanin. Phenoloxidase levels are widely used as an estimate of immunocompetence, especially against invading fungi and parasitoids (Rantala et al., 2000; Cerenius and Söderhäll, 2004; Barnes and Siva-Jothy, 2000; Mucklow and Ebert, 2003). However, studies in *Daphnia* reveal that pronounced genotypic variation in phenoloxidase activity complicates efforts to link host responses to immmunocompetence or immunopathology (Mucklow et al., 2004; Auld et al., 2013). Evaluating the functional consequences of these responses, therefore, requires accounting for other host defense mechanisms and pathogen traits.

With the two immune-mediated responses measured here, two different perspectives could help explain their functional effects. The first is that phenoloxidase and anorexia may help control pathogen populations (anti-growth resistance) and/or decrease the probability of infection (anti-infection resistance). For ‘A45’, anorexia and phenoloxidase were weakly and inversely related. Infection prevalence and spore yield were both minimal at the lower pathogen dose where food intake (and thus, pathogen exposure) increased slightly but phenoloxidase activity declined 9.5% from baseline. Infection prevalence and spore yields increased at the higher pathogen dose where food intake remained unchanged (relative to the controls) and phenoloxidase increased 28%. For ‘Midland 252’, anorexia and phenoloxidase were linearly related; phenoloxidase was lower when food intake (and pathogen exposure) were also low and vice versa. Low phenoloxidase and low food intake (and pathogen exposure) at lower spore doses resulted in higher infection rates relative to higher doses where both phenoloxidase and food intake (and pathogen exposure) increased. For ‘A45’ these mechanisms carried few fitness costs in terms of fecundity, growth, or survival. For ‘Midland 252’, fitness costs also remained relatively low, but the higher production of phenoloxidase (relative to ‘A45’) may explain that genotype’s slight decrease in reproduction.

An alternative perspective is that ‘A45’ was more susceptible at higher pathogen doses so the defense strategy was to decrease food intake and up-regulate phenoloxidase. Whereas ‘Midland 252’ was more susceptible at lower pathogen loads, thus this genotype hedged their bets by not decreasing food intake to offset the fitness costs of up-regulating phenoloxidase. The causality dilemma inherent in connecting these traits presents an interesting thought exercise that will hopefully catalyze future studies in systems with more molecular tools to tease apart these mechanisms. Irregardless of the cause and effect, these results join mounting evidence highlighting that beyond immune cells, defense against infectious agents involves an integrated and adaptively plastic arsenal of behavioral, physiological, and metabolic changes.

While the epidemiological and evolutionary importance of immunometabolism and associated changes in feeding behaviors are increasingly appreciated (Smith and Holt, 1996; Smith et al., 2015; Cotter et al., 2019; Pike et al., 2019), previous studies have largely focused on the molecular, physiological, or immunological underpinnings in single host genotypes (Adelman and Martin, 2009; Bashir-Tanoli and Tinsley, 2014; Adamo et al., 2007) or genetically-modified model systems (Schneider and Ayres, 2008; Rao et al., 2017). Further, these host-centric studies rarely track changes in host or pathogen life history traits (reviewed by Hite et al., 2020). As a consequence, the potential effects of immunometabolism and associated changes in feeding behaviors on host fitness and pathogen transmission remain poorly resolved.

We sought to answer two questions. Do more susceptible genotypes exhibit the strongest or weakest anorexia? Do the two focal host responses benefit the host, pathogen, both or neither? The answer is, at first glance, somewhat unsatisfactory: it depends on the host genotype and the environment. Once again, genotype x environment interactions matter. Yet, if we consider that ecology is founded on and propelled by the exciting goal of understanding the immense genetic diversity found in natural systems, these results reconfirm the need to move beyond non-model organisms to expand our understanding of immune polymorphism found in nature (Graham, 2021; Duffy et al., 2021). Such efforts require a greater appreciation for the fact the defense strategies fall along a spectrum that cannot be neatly categorized as resistance or tolerance. Of course, these efforts will require the development of molecular tools for non-model organisms. We do not underestimate the logistical challenges associated with such endeavors, but rather join recent studies emphasizing the need for these tools and collaborations. In the meantime, approaches such as ours that combine tools from population ecology and immunology provide valuable insight on how natural selection shapes strategies for host defense and pathogen transmission.

## Acknowledgements

We are grateful to Spencer Hall and Meghan Duffy who generously shared host genotypes and pathogen spores. We thank the Duffy lab and Daniel Medina for helpful discussions and feedback on the manuscript. Dirk Anderson at the UNL Flow Cytometry Core Facility provided invaluable assistance with flow cytometry. Funding was provided by the National Institutes of Health (F32GM12846) to JLH.

## Author’s Contributions

J.L.H. designed the study and analysed the data; J.L.H, A.C.P.B, and R.E.V. conducted the experiment and wrote the manuscript. All authors contributed to revisions.

## Data Availability Statement

All data and R code are publicly available from the GitHub repository (https://github.com/alainapb/Tolerance_Resistance).

