## Appendix for "Natural variation in host defense strategies impacts both host and pathogen fitness"

APB

7/23/2022

#### Comparison of spore counts using flow cytometry to traditional counts using a hemocytometer in 'Standard' genotype

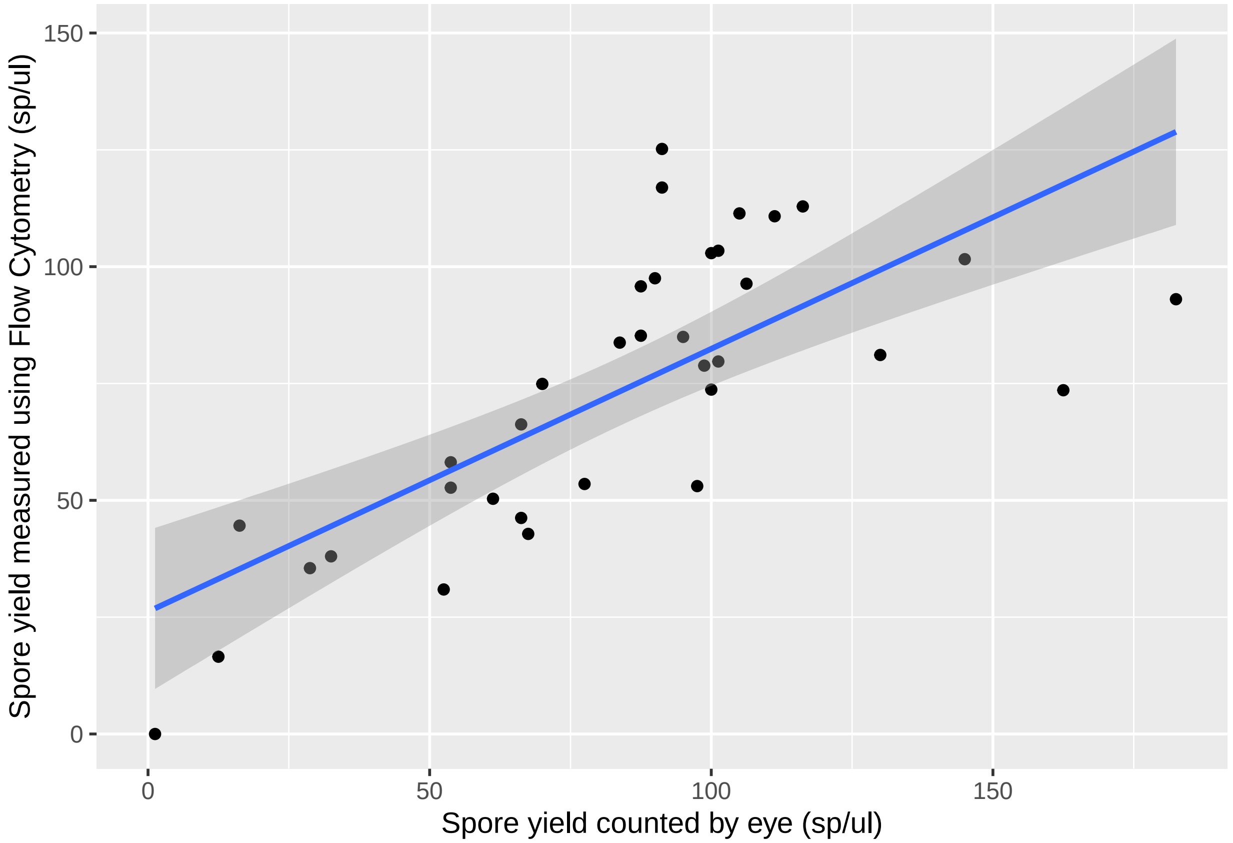


```r
fit<-lm(sp.ul ~ flow_sp_ul - 1, data = sporecheck)
# R squared
summary(fit)$r.squared
```

```
## [1] 0.9130979
```

**Survival Analyses**

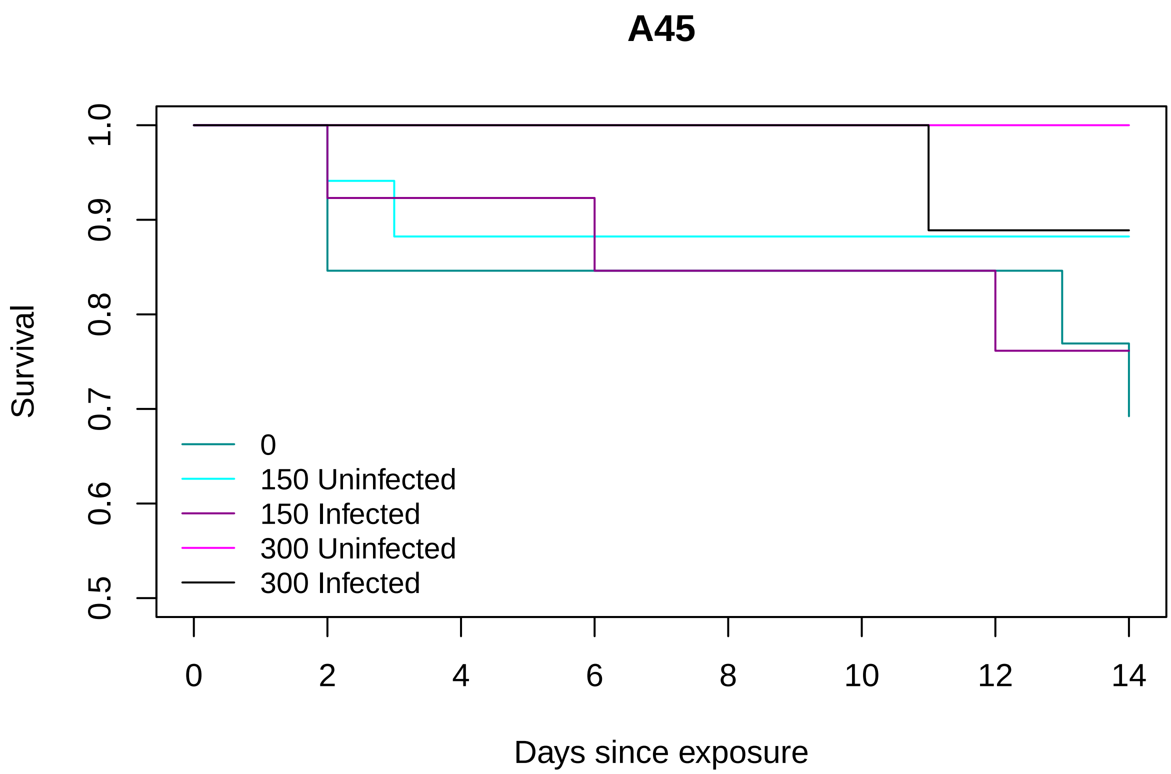


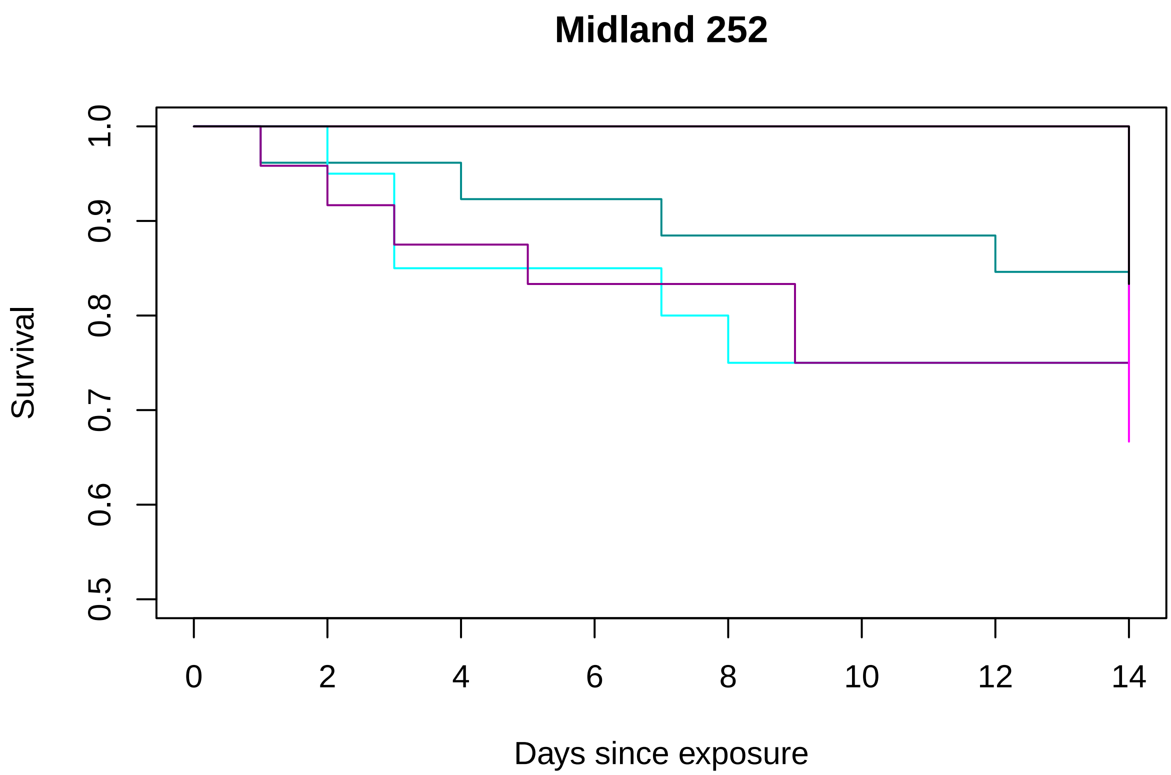

#### Survival Analyses Continued

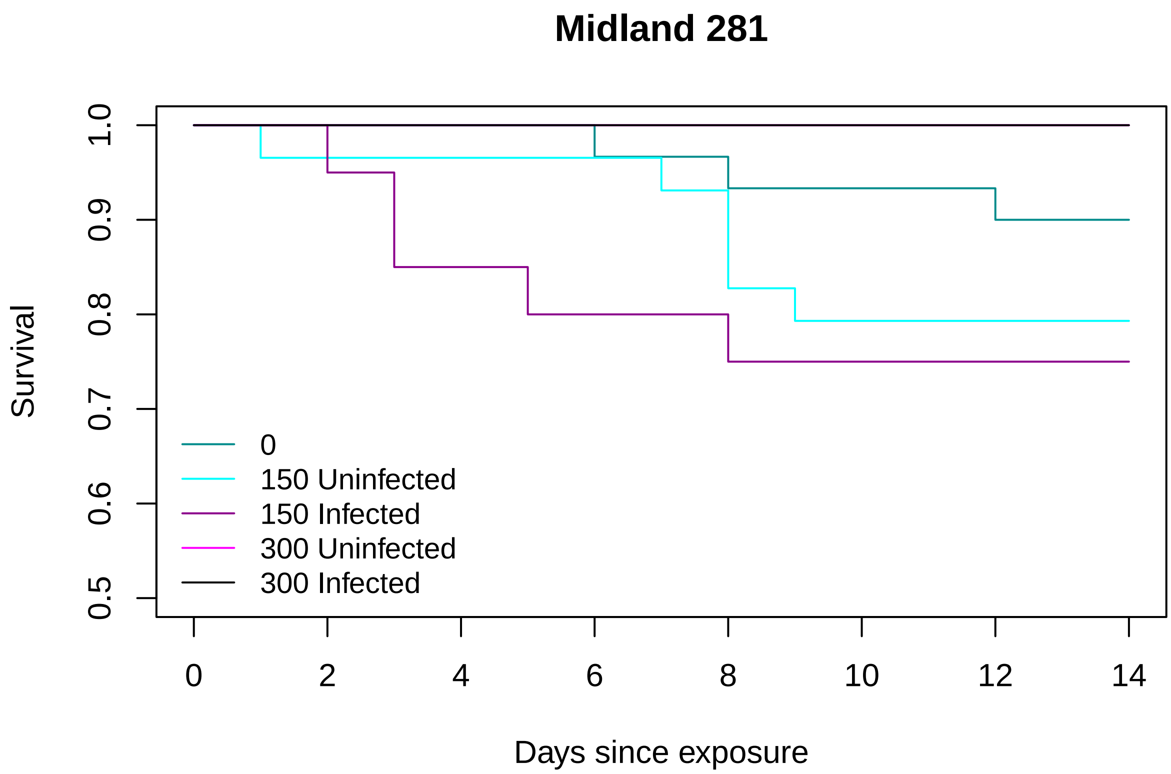


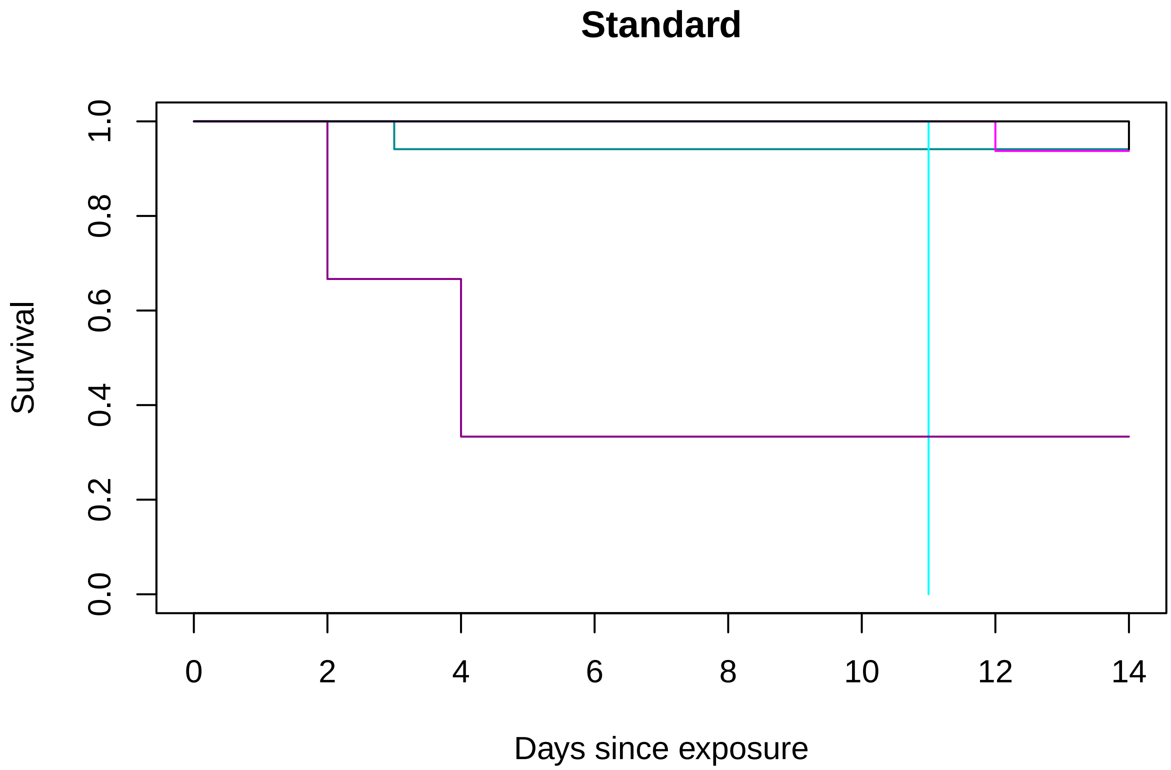

In Standard, there was only one individual that was exposed to 150 sp/ml that did not get infected and that individual died eleven days after the exposure (blue line).

#### Impacts of Feeding Behavior

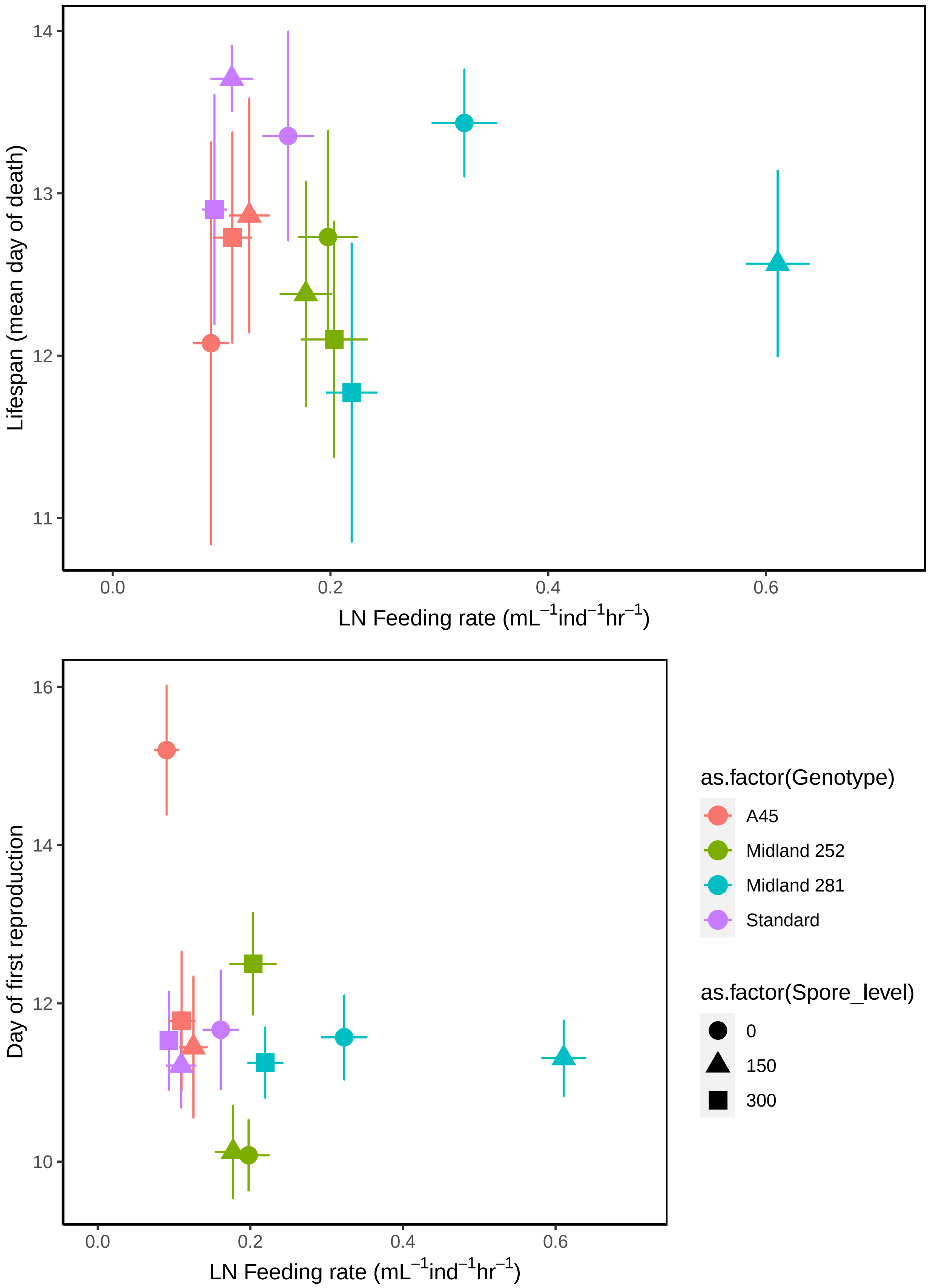

#### Table 2

```
##
## Call:
## glm(formula = mean_feeding_rate ~ Genotype * Size_mm2 + Infected_Uninfected +
##     Spore_level, family = gaussian(link = "log"), data = total)
##
## Deviance Residuals:
##      Min        1Q    Median        3Q       Max
## -0.28232  -0.07199  -0.00455   0.07947   0.63002
##
## Coefficients:
##                               Estimate Std. Error t value Pr(>|t|)
## (Intercept)                 -3.6790349  1.3734430  -2.679  0.00786 **
## GenotypeMidland 252          0.4129904  1.4218075   0.290  0.77169
## GenotypeMidland 281          1.7544277  1.3921475   1.260  0.20870
## GenotypeStandard             0.8783600  1.8305967   0.480  0.63175
## Size_mm2                     0.8337302  0.8508144   0.980  0.32803
## Infected_UninfectedExposed   1.1843364  0.1265402   9.359  < 2e-16 ***
## Infected_UninfectedInfected  0.9761503  0.1852444   5.270 2.85e-07 ***
## Spore_level                 -0.0046333  0.0006209  -7.462 1.25e-12 ***
## GenotypeMidland 252:Size_mm2  0.1153302  0.8790541   0.131  0.89572
## GenotypeMidland 281:Size_mm2 -0.2638278  0.8628215  -0.306  0.76002
## GenotypeStandard:Size_mm2   -0.4227152  1.1498522  -0.368  0.71345
## ---
## Signif. codes:  0 '***' 0.001 '**' 0.01 '*' 0.05 '.' 0.1 ' ' 1
##
## (Dispersion parameter for gaussian family taken to be 0.01640027)
##
##     Null deviance: 10.296  on 273  degrees of freedom
## Residual deviance:  4.313  on 263  degrees of freedom
##   (7 observations deleted due to missingness)
## AIC: -335.93
##
## Number of Fisher Scoring iterations: 9

## Analysis of Deviance Table (Type III tests)
##
## Response: mean_feeding_rate
##                     LR Chisq Df Pr(>Chisq)
## Genotype               9.269  3    0.02592 *
## Size_mm2               0.789  1    0.37434
## Infected_Uninfected   91.087  2    < 2e-16 ***
## Spore_level           77.146  1    < 2e-16 ***
## Genotype:Size_mm2      1.833  3    0.60772
## ---
## Signif. codes:  0 '***' 0.001 '**' 0.01 '*' 0.05 '.' 0.1 ' ' 1

## Analysis of Deviance Table (Type II tests)
##
## Response: mean_feeding_rate
##                     LR Chisq Df Pr(>Chisq)
## Genotype             218.487  3  < 2.2e-16 ***
```

```
## Size_mm2              29.557  1  5.429e-08 ***
## Infected_Uninfected   91.087  2  < 2.2e-16 ***
## Spore_level           77.146  1  < 2.2e-16 ***
## Genotype:Size_mm2      1.833  3     0.6077
## ---
## Signif. codes:  0 '***' 0.001 '**' 0.01 '*' 0.05 '.' 0.1 ' ' 1
```

```
##       dAICc df
## mod1c  0.0  9
## mod1b  4.6  12
```

```
##
## Call:
## glm(formula = mean_feeding_rate ~ Genotype + Size_mm2 + Infected_Uninfected +
##     Spore_level, family = gaussian(link = "log"), data = total)
##
## Deviance Residuals:
##      Min        1Q    Median        3Q       Max
## -0.28940  -0.07688  -0.00773   0.07413   0.62273
##
## Coefficients:
##                              Estimate Std. Error t value Pr(>|t|)
## (Intercept)                -3.3847228  0.2514575 -13.460  < 2e-16 ***
## GenotypeMidland 252         0.5905779  0.1726311   3.421 0.000722 ***
## GenotypeMidland 281         1.3315642  0.1602593   8.309 4.99e-15 ***
## GenotypeStandard            0.2143465  0.2240725   0.957 0.339640
## Size_mm2                    0.6532189  0.1179270   5.539 7.30e-08 ***
## Infected_UninfectedExposed   1.1776463  0.1260141   9.345  < 2e-16 ***
## Infected_UninfectedInfected  0.9654600  0.1846276   5.229 3.45e-07 ***
## Spore_level                -0.0046315  0.0006148  -7.534 7.73e-13 ***
## ---
## Signif. codes:  0 '***' 0.001 '**' 0.01 '*' 0.05 '.' 0.1 ' ' 1
##
## (Dispersion parameter for gaussian family taken to be 0.01632856)
##
##     Null deviance: 10.2961  on 273  degrees of freedom
## Residual deviance:  4.3431  on 266  degrees of freedom
##   (7 observations deleted due to missingness)
## AIC: -340.03
##
## Number of Fisher Scoring iterations: 8
```

| Term | LR Chisq | Df | P |
|---|---|---|---|
| Genotype | 219.44633 | 3 | 0.00000000000000000000000000000000000000000000 |
| Size_mm2 | 29.68707 | 1 | 0.00000005077239218380875567536151984730785624 |
| Infected_Uninfected | 90.68973 | 2 | 0.0000000000000000000202756924730664848386479 |
| Spore_level | 78.06434 | 1 | 0.00000000000000000099738420801168999133819181 |

```
## Genotype = A45:
```

```
##  contrast           estimate    SE  df t.ratio p.value
##  Control - Exposed    -1.178 0.126 266  -9.345  <.0001
##  Control - Infected   -0.965 0.185 266  -5.229  <.0001
##  Exposed - Infected    0.212 0.140 266   1.512  0.2867
##
## Genotype = Midland 252:
##  contrast           estimate    SE  df t.ratio p.value
##  Control - Exposed    -1.178 0.126 266  -9.345  <.0001
##  Control - Infected   -0.965 0.185 266  -5.229  <.0001
##  Exposed - Infected    0.212 0.140 266   1.512  0.2867
##
## Genotype = Midland 281:
##  contrast           estimate    SE  df t.ratio p.value
##  Control - Exposed    -1.178 0.126 266  -9.345  <.0001
##  Control - Infected   -0.965 0.185 266  -5.229  <.0001
##  Exposed - Infected    0.212 0.140 266   1.512  0.2867
##
## Genotype = Standard:
##  contrast           estimate    SE  df t.ratio p.value
##  Control - Exposed    -1.178 0.126 266  -9.345  <.0001
##  Control - Infected   -0.965 0.185 266  -5.229  <.0001
##  Exposed - Infected    0.212 0.140 266   1.512  0.2867
##
## Results are given on the log (not the response) scale.
## P value adjustment: tukey method for comparing a family of 3 estimates
```

### with size correction

```
## Analysis of Deviance Table (Type III tests)
##
## Response: meanfeedrate
##                   LR Chisq Df Pr(>Chisq)
## probinf              3.527  1    0.06038 .
## finalsize            2.749  1    0.09729 .
## Spore_level          0.964  1    0.32623
## Genotype           105.966  3    < 2e-16 ***
## probinf:finalsize    3.566  1    0.05898 .
## ---
## Signif. codes:  0 '***' 0.001 '**' 0.01 '*' 0.05 '.' 0.1 ' ' 1
```

```
## Analysis of Deviance Table (Type III tests)
##
## Response: meanfeedrate
##                   LR Chisq Df Pr(>Chisq)
## probinf              3.230  1    0.07232 .
## finalsize            2.748  1    0.09737 .
## Genotype           105.663  3    < 2e-16 ***
## probinf:finalsize    3.466  1    0.06263 .
## ---
## Signif. codes:  0 '***' 0.001 '**' 0.01 '*' 0.05 '.' 0.1 ' ' 1
```

```
## Analysis of Deviance Table (Type III tests)
```

```
##
## Response: meanfeedrate
##                   LR Chisq Df Pr(>Chisq)
## probinf              3.527  1    0.06038 .
## finalsize            2.749  1    0.09729 .
## Spore_level          0.964  1    0.32623
## Genotype           105.966  3    < 2e-16 ***
## probinf:finalsize    3.566  1    0.05898 .
## ---
## Signif. codes:  0 '***' 0.001 '**' 0.01 '*' 0.05 '.' 0.1 ' ' 1
```

```
## Analysis of Deviance Table (Type III tests)
##
## Response: meanfeedrate
##                   LR Chisq Df Pr(>Chisq)
## probinf              3.081  1    0.07920 .
## finalsize            1.369  1    0.24192
## Genotype           105.706  3    < 2e-16 ***
## probinf:finalsize    3.384  1    0.06583 .
## ---
## Signif. codes:  0 '***' 0.001 '**' 0.01 '*' 0.05 '.' 0.1 ' ' 1
```

```
##             dAICc df
## modfeedinfb 0.0   8
## modfeedinfd 0.4   8
## modfeedinf  1.2   9
## modfeedinfc 1.2   9
```

```
##
## Call:
## glm(formula = meanfeedrate ~ probinf * finalsize + Genotype,
##     family = gaussian(link = "identity"), data = inf1)
##
## Deviance Residuals:
##      Min        1Q    Median        3Q       Max
## -0.39125  -0.08163  -0.01700   0.07431   0.53012
##
## Coefficients:
##                     Estimate Std. Error t value Pr(>|t|)
## (Intercept)         -0.17391    0.17409  -0.999  0.31894
## probinf              0.37294    0.20752   1.797  0.07373 .
## finalsize            0.10804    0.06518   1.658  0.09884 .
## GenotypeMidland 252  0.11927    0.03774   3.160  0.00181 **
## GenotypeMidland 281  0.30358    0.03335   9.102  < 2e-16 ***
## GenotypeStandard     0.03748    0.03485   1.075  0.28337
## probinf:finalsize   -0.15923    0.08552  -1.862  0.06401 .
## ---
## Signif. codes:  0 '***' 0.001 '**' 0.01 '*' 0.05 '.' 0.1 ' ' 1
##
## (Dispersion parameter for gaussian family taken to be 0.02436518)
##
##     Null deviance: 8.3950  on 220  degrees of freedom
## Residual deviance: 5.2141  on 214  degrees of freedom
```

```
##   (60 observations deleted due to missingness)
## AIC: -184.87
##
## Number of Fisher Scoring iterations: 2
```

```
## Analysis of Deviance Table (Type III tests)
##
## Response: meanfeedrate
##                   LR Chisq Df Pr(>Chisq)
## probinf              3.230  1    0.07232 .
## finalsize            2.748  1    0.09737 .
## Genotype           105.663  3    < 2e-16 ***
## probinf:finalsize    3.466  1    0.06263 .
## ---
## Signif. codes:  0 '***' 0.001 '**' 0.01 '*' 0.05 '.' 0.1 ' ' 1
```

| Term | LR Chisq | Df | Pr(>Chisq) |
|---|---:|:---:|---:|
| probinf | 3.229664 | 1 | 0.07231552365034664420218035730 |
| finalsize | 2.748078 | 1 | 0.09737143128251564416775210020 |
| Genotype | 105.662556 | 3 | 0.00000000000000000000009410693 |
| probinf:finalsize | 3.466209 | 1 | 0.06263468621257345381181380617 |

#### Impacts of Infection

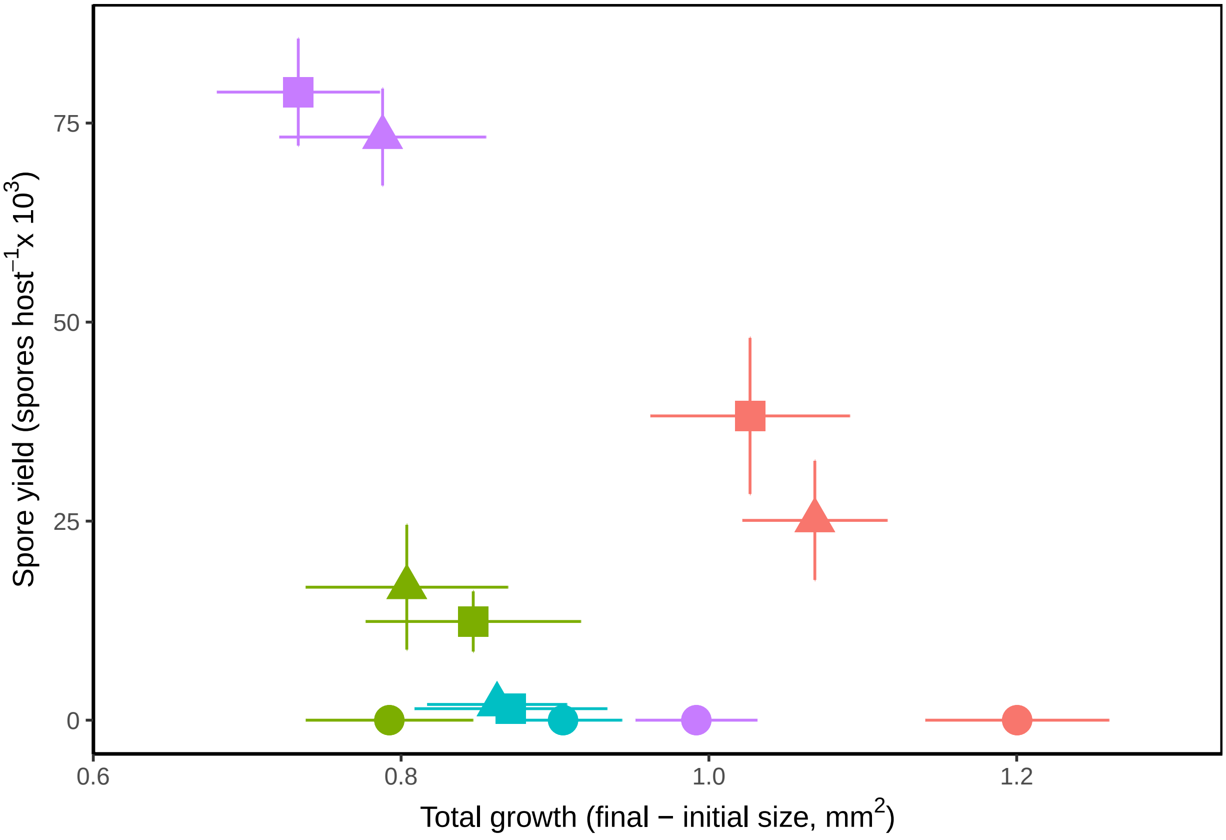


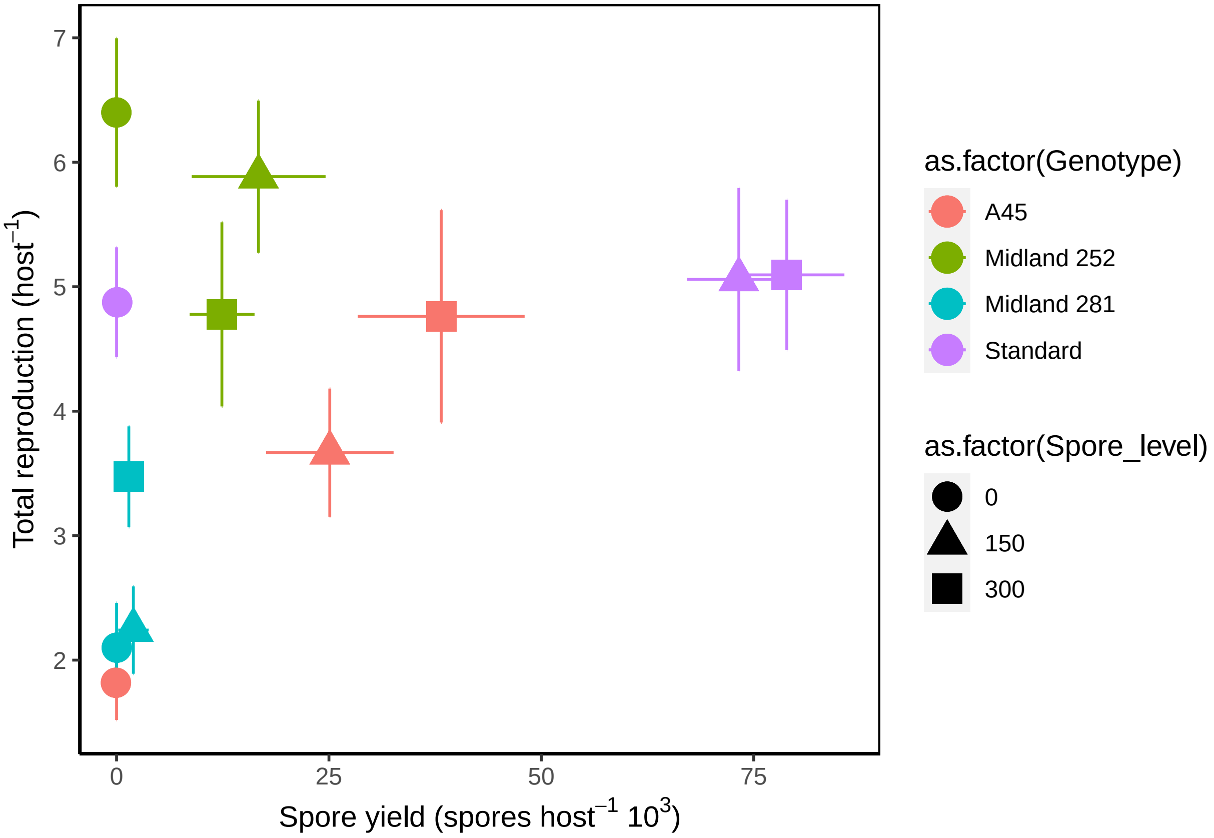

### Table 5a

```
##       dAICc df
## mod3m 0     3
## mod2m 1     5
```

```
##
## Call:
## glm(formula = y ~ mean_feeding_rate + Size_mm2, family = binomial,
##     data = infected)
##
## Deviance Residuals:
##     Min       1Q   Median       3Q      Max
## -1.0168  -0.7848  -0.5477   0.4820   1.8138
##
## Coefficients:
##                   Estimate Std. Error z value Pr(>|z|)
## (Intercept)        -1.4615     0.8171  -1.789  0.07368 .
## mean_feeding_rate  -3.2139     1.0594  -3.034  0.00242 **
## Size_mm2            0.6494     0.5396   1.204  0.22877
## ---
## Signif. codes:  0 '***' 0.001 '**' 0.01 '*' 0.05 '.' 0.1 ' ' 1
##
## (Dispersion parameter for binomial family taken to be 1)
##
##     Null deviance: 108.639  on 188  degrees of freedom
## Residual deviance:  95.683  on 186  degrees of freedom
##   (6 observations deleted due to missingness)
## AIC: 193.18
##
## Number of Fisher Scoring iterations: 5
```

```
## Analysis of Deviance Table (Type III tests)
##
## Response: y
##                   LR Chisq Df Pr(>Chisq)
## mean_feeding_rate   12.272  1  0.0004597 ***
## Size_mm2             1.494  1  0.2215954
## ---
## Signif. codes:  0 '***' 0.001 '**' 0.01 '*' 0.05 '.' 0.1 ' ' 1
```

| Term | LR Chisq | Df | Pr(>Chisq) |
|---|---|---|---|
| mean_feeding_rate | 12.27225 | 1 | 0.000459744 |
| Size_mm2 | 1.49401 | 1 | 0.221595370 |

### Table 5b (accounting for genetic variation)

```
## Analysis of Deviance Table (Type II tests)
##
```

```
## Response: y
##                           Df   Chisq Pr(>Chisq)
## mean_feeding_rate          1  0.0219  0.8823744
## Size_mm2                   1  0.3861  0.5343357
## as.factor(Genotype)        3 16.7564  0.0007931 ***
## as.factor(Spore_level)     1  0.1030  0.7482329
## mean_feeding_rate:Size_mm2  1  3.6375  0.0564906 .
## ---
## Signif. codes:  0 '***' 0.001 '**' 0.01 '*' 0.05 '.' 0.1 ' ' 1

## Analysis of Deviance Table (Type II tests)
##
## Response: y
##                       Df   Chisq Pr(>Chisq)
## mean_feeding_rate      1  0.1860  0.6662878
## Size_mm2               1  0.5353  0.4643890
## as.factor(Genotype)    3 16.6723  0.0008253 ***
## as.factor(Spore_level)  1  0.0310  0.8602011
## ---
## Signif. codes:  0 '***' 0.001 '**' 0.01 '*' 0.05 '.' 0.1 ' ' 1

##       dAICc df
## mod2h 0.0   8
## mod2i 1.4   7

## Analysis of Deviance Table (Type III tests)
##
## Response: y
##                           LR Chisq Df Pr(>Chisq)
## mean_feeding_rate            3.1284  1  0.0769370 .
## Size_mm2                     3.6215  1  0.0570377 .
## as.factor(Genotype)         19.3611  3  0.0002302 ***
## as.factor(Spore_level)       0.1031  1  0.7481421
## mean_feeding_rate:Size_mm2   3.5805  1  0.0584600 .
## ---
## Signif. codes:  0 '***' 0.001 '**' 0.01 '*' 0.05 '.' 0.1 ' ' 1

## Analysis of Deviance Table (Type III tests)
##
## Response: y
##                           LR Chisq Df Pr(>Chisq)
## mean_feeding_rate            3.1284  1  0.0769370 .
## Size_mm2                     3.6215  1  0.0570377 .
## as.factor(Genotype)         19.3611  3  0.0002302 ***
## as.factor(Spore_level)       0.1031  1  0.7481421
## mean_feeding_rate:Size_mm2   3.5805  1  0.0584600 .
## ---
## Signif. codes:  0 '***' 0.001 '**' 0.01 '*' 0.05 '.' 0.1 ' ' 1
```

| Term | LR Chisq | Df | Pr(>Chisq) |
|---|---|---|---|
| mean_feeding_rate | 3.1284463 | 1 | 0.0769370344 |
| Size_mm2 | 3.6214925 | 1 | 0.0570376833 |
| as.factor(Genotype) | 19.3610810 | 3 | 0.0002302002 |
| as.factor(Spore_level) | 0.1030989 | 1 | 0.7481420645 |
| mean_feeding_rate:Size_mm2 | 3.5805454 | 1 | 0.0584599561 |

### Table 5a Spore yield

```
##
## Call:
## glm(formula = Spores_ul ~ mean_feeding_rate + as.factor(Spore_level),
##     data = total)
##
## Deviance Residuals:
##     Min       1Q   Median       3Q      Max
## -39.925  -24.637   -3.703   11.404  138.041
##
## Coefficients:
##                           Estimate Std. Error t value Pr(>|t|)
## (Intercept)                 10.216      4.128   2.475    0.014 *
## mean_feeding_rate          -46.881     10.760  -4.357 1.95e-05 ***
## as.factor(Spore_level)150   27.941      4.932   5.665 4.20e-08 ***
## as.factor(Spore_level)300   30.621      4.991   6.135 3.49e-09 ***
## ---
## Signif. codes:  0 '***' 0.001 '**' 0.01 '*' 0.05 '.' 0.1 ' ' 1
##
## (Dispersion parameter for gaussian family taken to be 992.8513)
##
##     Null deviance: 307338  on 243  degrees of freedom
## Residual deviance: 238284  on 240  degrees of freedom
##   (37 observations deleted due to missingness)
## AIC: 2382.2
##
## Number of Fisher Scoring iterations: 2
```

```
## Analysis of Deviance Table (Type III tests)
##
## Response: Spores_ul
##                        LR Chisq Df Pr(>Chisq)
## mean_feeding_rate        18.984  1  1.318e-05 ***
## as.factor(Spore_level)   48.256  2  3.321e-11 ***
## ---
## Signif. codes:  0 '***' 0.001 '**' 0.01 '*' 0.05 '.' 0.1 ' ' 1
```

```
## Call:
##    aov(formula = mod2k)
##
```

```
## Terms:
##                 mean_feeding_rate as.factor(Spore_level) Residuals
## Sum of Squares           21142.83               47911.20 238284.30
## Deg. of Freedom                 1                      2       240
##
## Residual standard error: 31.50954
## Estimated effects may be unbalanced
## 37 observations deleted due to missingness
```

```
## Analysis of Deviance Table (Type III tests)
##
## Response: Spores_ul
##                            LR Chisq Df Pr(>Chisq)
## mean_feeding_rate           0.22312  1     0.6367
## Size_mm2                    2.19853  1     0.1381
## Spore_level_fct             0.06526  1     0.7984
## mean_feeding_rate:Size_mm2  0.65240  1     0.4193
```

```
## Analysis of Deviance Table (Type III tests)
##
## Response: Spores_ul
##                                LR Chisq Df Pr(>Chisq)
## mean_feeding_rate               0.79702  1     0.3720
## Size_mm2                        0.08375  1     0.7723
## Spore_level_fct                 1.05072  1     0.3053
## Final_Size_mm2                  0.55722  1     0.4554
## Spore_level_fct:Final_Size_mm2  1.28394  1     0.2572
```

```
## Analysis of Deviance Table (Type III tests)
##
## Response: Spores_ul
##                 LR Chisq Df Pr(>Chisq)
## Spore_level_fct   0.4621  1     0.4967
## Final_Size_mm2    5.2617  1     0.0218 *
## ---
## Signif. codes:  0 '***' 0.001 '**' 0.01 '*' 0.05 '.' 0.1 ' ' 1
```

```
##                df      AIC
## modfeedspores1  6 644.8335
## modfeedspores2  7 610.9386
## modfeedspores3  6 610.3441
## modfeedspores4  4 607.3463
```

### Table 5b Spore yield

```
## Analysis of Deviance Table (Type III tests)
##
## Response: Spores_ul
##                            LR Chisq Df Pr(>Chisq)
## mean_feeding_rate            1.4557  1    0.22762
## Size_mm2                     0.0266  1    0.87046
## Spore_level_fct              1.0240  1    0.31158
```

```
## Final_Size_mm2               6.6193  1    0.01009 *
## Genotype                     6.7155  3    0.08154 .
## mean_feeding_rate:Size_mm2   1.4536  1    0.22796
## ---
## Signif. codes:  0 '***' 0.001 '**' 0.01 '*' 0.05 '.' 0.1 ' ' 1

## Analysis of Deviance Table (Type III tests)
##
## Response: Spores_ul
##                   LR Chisq Df Pr(>Chisq)
## mean_feeding_rate   0.0253  1    0.87353
## Size_mm2            0.8198  1    0.36524
## Spore_level_fct     0.7896  1    0.37421
## Final_Size_mm2      6.2452  1    0.01245 *
## Genotype            5.6513  3    0.12986
## ---
## Signif. codes:  0 '***' 0.001 '**' 0.01 '*' 0.05 '.' 0.1 ' ' 1

## Analysis of Deviance Table (Type III tests)
##
## Response: Spores_ul
##                   LR Chisq Df Pr(>Chisq)
## mean_feeding_rate   0.2142  1     0.6435
## Size_mm2            0.6809  1     0.4093
## Spore_level_fct     0.0722  1     0.7881
## Genotype            3.2987  3     0.3478

## Analysis of Deviance Table (Type III tests)
##
## Response: Spores_ul
##                   LR Chisq Df Pr(>Chisq)
## mean_feeding_rate    1.214  1     0.2705
## Spore_level_fct     52.335  2  4.320e-12 ***
## Genotype            74.429  3  4.804e-16 ***
## ---
## Signif. codes:  0 '***' 0.001 '**' 0.01 '*' 0.05 '.' 0.1 ' ' 1

## Analysis of Deviance Table (Type III tests)
##
## Response: Spores_ul
##                 LR Chisq Df Pr(>Chisq)
## Spore_level_fct   51.567  2  6.343e-12 ***
## Genotype          97.760  3  < 2.2e-16 ***
## ---
## Signif. codes:  0 '***' 0.001 '**' 0.01 '*' 0.05 '.' 0.1 ' ' 1

##
## Call:
## glm(formula = Spores_ul ~ mean_feeding_rate + Size_mm2 + Spore_level_fct +
##     Final_Size_mm2 + Genotype, family = gaussian(link = "identity"),
##     data = totalpos)
##
## Deviance Residuals:
```

```
##     Min       1Q   Median       3Q      Max
## -52.833  -23.933   -0.319   20.326   95.758
##
## Coefficients:
##                     Estimate Std. Error t value Pr(>|t|)
## (Intercept)          -12.719     37.399  -0.340   0.7351
## mean_feeding_rate      5.891     37.012   0.159   0.8741
## Size_mm2             -16.291     17.993  -0.905   0.3693
## Spore_level_fct300     7.395      8.322   0.889   0.3782
## Final_Size_mm2        40.740     16.302   2.499   0.0155 *
## GenotypeMidland 252    5.831     14.571   0.400   0.6906
## GenotypeMidland 281  -27.677     26.048  -1.063   0.2927
## GenotypeStandard      18.712     10.329   1.811   0.0756 .
## ---
## Signif. codes:  0 '***' 0.001 '**' 0.01 '*' 0.05 '.' 0.1 ' ' 1
##
## (Dispersion parameter for gaussian family taken to be 945.2518)
##
##     Null deviance: 62483  on 61  degrees of freedom
## Residual deviance: 51044  on 54  degrees of freedom
##   (3 observations deleted due to missingness)
## AIC: 610.17
##
## Number of Fisher Scoring iterations: 2
```

```
## Analysis of Deviance Table (Type III tests)
##
## Response: Spores_ul
##                   LR Chisq Df Pr(>Chisq)
## mean_feeding_rate   0.0253  1    0.87353
## Size_mm2            0.8198  1    0.36524
## Spore_level_fct     0.7896  1    0.37421
## Final_Size_mm2      6.2452  1    0.01245 *
## Genotype            5.6513  3    0.12986
## ---
## Signif. codes:  0 '***' 0.001 '**' 0.01 '*' 0.05 '.' 0.1 ' ' 1
```

| Term | LR Chisq | Df | Pr(>Chisq) |
|---|---|---|---|
| mean_feeding_rate | 0.02533612 | 1 | 0.87353240 |
| Size_mm2 | 0.81979692 | 1 | 0.36523955 |
| Spore_level_fct | 0.78963588 | 1 | 0.37421025 |
| Final_Size_mm2 | 6.24521611 | 1 | 0.01245292 |
| Genotype | 5.65134086 | 3 | 0.12986181 |

```
## Spore_level_fct = 150:
##  contrast                  estimate   SE df t.ratio p.value
##  A45 - Midland 252            -5.83 14.6 54  -0.400  0.9781
##  A45 - Midland 281            27.68 26.0 54   1.063  0.7135
##  A45 - Standard              -18.71 10.3 54  -1.811  0.2792
```

```
##  Midland 252 - Midland 281    33.51 25.2 54   1.327  0.5500
##  Midland 252 - Standard      -12.88 12.6 54  -1.024  0.7363
##  Midland 281 - Standard      -46.39 25.8 54  -1.801  0.2840
##
## Spore_level_fct = 300:
##  contrast                  estimate   SE df t.ratio p.value
##  A45 - Midland 252            -5.83 14.6 54  -0.400  0.9781
##  A45 - Midland 281            27.68 26.0 54   1.063  0.7135
##  A45 - Standard              -18.71 10.3 54  -1.811  0.2792
##  Midland 252 - Midland 281    33.51 25.2 54   1.327  0.5500
##  Midland 252 - Standard      -12.88 12.6 54  -1.024  0.7363
##  Midland 281 - Standard      -46.39 25.8 54  -1.801  0.2840
##
## P value adjustment: tukey method for comparing a family of 4 estimates
```
